# Bacterial metabolites induce cell wall remodeling, antifungal resistance, and immune recognition of commensal fungi

**DOI:** 10.1101/2025.07.26.666966

**Authors:** Faith Anderson Davis, Kalpana Singh, Joseph M. Krampen, Jaidyn A Bryant, Kyla S. Ost, Shannon E. Righi, Marcy J. Balunas, Tuo Wang, Teresa R. O’Meara

## Abstract

The fungus *Candida albicans* commensally colonizes mucosal surfaces in healthy individuals but can cause both superficial mucosal and life-threatening disseminated infections. The balance between commensalism and pathogenicity is complex and depends on factors including host and fungal genetic background, the host environment, and fungal interactions with local microbes. The major interaction interface of *C. albicans* with the host is its multilayered cell wall, which is dynamic and highly responsive to the surrounding environment. Therefore, factors that influence the fungal cell wall will directly impact *C. albicans*-host interactions. Our work demonstrates that multiple physiologically-relevant gastrointestinal bacteria influence fungal cell wall composition during co-culture with *C. albicans,* including as complex communities derived from the gut. Using *Escherichia coli* as a model, we show that bacterial-induced fungal cell wall remodeling occurs rapidly and is mediated by secreted bacterial metabolite(s). Fungal mutant analysis revealed that the high osmolarity glycerol (HOG) pathway, which is critical for responding to environmental stresses, has an important role in regulating this cell wall remodeling phenotype through the Sln1 histidine kinase. Importantly, bacterial-mediated fungal cell wall remodeling increases *C. albicans* resistance to the echinocandins, increases recognition by both dectin-1 and dectin-2, and decreases recognition by human IgA. Overall, this work comprehensively characterizes an interaction between *C. albicans* and common gastrointestinal bacteria that has important implications for fungal biology and host interactions.

## INTRODUCTION

*Candida albicans* can exist as a commensal member of the mucosal microbiota, but also has the capacity to cause a range of disease, from superficial mucosal infections to life-threatening disseminated infections ^1^. The balance between commensalism and pathogenicity is complex and multifactorial, depending on the genetic backgrounds of both host and fungus, host immune status, and fungal interactions with the local microbiota ^2–6^, all of which can be influenced by *C. albicans* morphological transitions, metabolic changes, and cell wall remodeling ^7,8^. As the main interface between *C. albicans* and the surrounding environment, the fungal cell wall contributes to structural integrity, sensing and responding to environmental stress, and mediating interactions with host cells. Cell wall composition is highly responsive to changes in environmental conditions, including nutrient source, oxygen content, metal availability, pH, or temperature ^9–15^, and these changes impact antifungal sensitivity, tolerance to stress, and importantly, recognition by host immune cells by altering PAMP exposure ^16–21^.

Recently, we demonstrated that human commensal *C. albicans* isolates retain the capacity to filament and cause disease ^22^. Although experimental evolution of *C. albicans* SC5314 through the murine gastrointestinal tract consistently produced yeast-locked strains with higher competitive fitness and decreased virulence, this outcome only occurred in mice maintained on antibiotics ^23^. Moreover, *C. albicans* exists primarily in the yeast form during gastrointestinal colonization of germ-free mice, but displays a mixture of yeast and hyphal morphologies during colonization of conventionally raised, antibiotic-treated mice ^24^. Together, this implies that an intact microbiota is important for applying selective pressure on *C. albicans* to maintain hyphal morphogenesis and virulence programs. However, the mechanisms at this inter-microbial interface are still being explored.

Bacteria have profound impacts on fungal biology, colonization, and virulence, both *in vitro* and *in vivo*. Commensal anaerobic bacteria induce production of the antimicrobial peptide CRAMP (LL-37 in humans), which reduces *C. albicans* gastrointestinal colonization and virulence potential, and bacterial-derived secondary metabolites, such as short chain fatty acids (SCFAs), alter *C. albicans* growth, colonization levels, morphology, and cell wall composition ^5,25–29^. Several studies have identified bacterial compounds that influence *C. albicans* biology by inhibiting filamentation, inhibiting growth, or directly killing fungal cells ^30–34^. However, the interplay between bacteria and the fungal cell wall has only recently begun to be explored. *Bacteroides* species can degrade purified fungal cell wall components, including mannan and β-1,6-glucan ^35,36^, but the impact of this during co-colonization is unclear. Recent work has shown evidence that co-incubation with live bacteria or bacterial supernatants can change fungal cell wall composition, although it is not clear how bacteria alter the fungal cell wall or how these changes impact fungal immune recognition^28,37^.

Here, we demonstrate that bacterial species induce fungal cell wall remodeling, with *E. coli* having the most significant impact on *C. albicans* cell wall composition. In contrast to cell wall masking models, we observe increased exposure of both β-glucan and mannan after co-incubation with bacteria, resulting in increased immunogenicity. We show that secreted bacterial metabolite(s) are responsible for inducing the remodeling phenotype, and a central stress pathway in *C. albicans* is required for the full remodeling response. Critically, we observed that bacterial-mediated fungal cell wall remodeling alters antifungal susceptibility and both innate and adaptive immune responses to *C. albicans*. These results underscore the importance of investigating fungal-bacterial interactions and how they may alter fungal virulence and disease outcomes.

## Results

### *Candida albicans* undergoes global cell wall remodeling during co-culture with *Escherichia coli*

To investigate the impact of common gastrointestinal-colonizing bacteria on the *Candida albicans* cell wall, we first developed an *in vitro* co-culture model. We grew *C. albicans* strain SC5314 for 24 hours in rich media, Yeast Peptone Dextrose (YPD), in monoculture or co-culture with *Escherichia coli* strain MC1061. The standard growth conditions for *C. albicans*, YPD media at 30°C, also allows for growth of *E. coli* (SFig 1). Although we observed a decrease in *C. albicans* growth rate during co-culture with *E. coli*, fungal cells remained viable following 24 hours of co-culture, based on propidium iodide staining (SFig 2).

**Figure 1:**
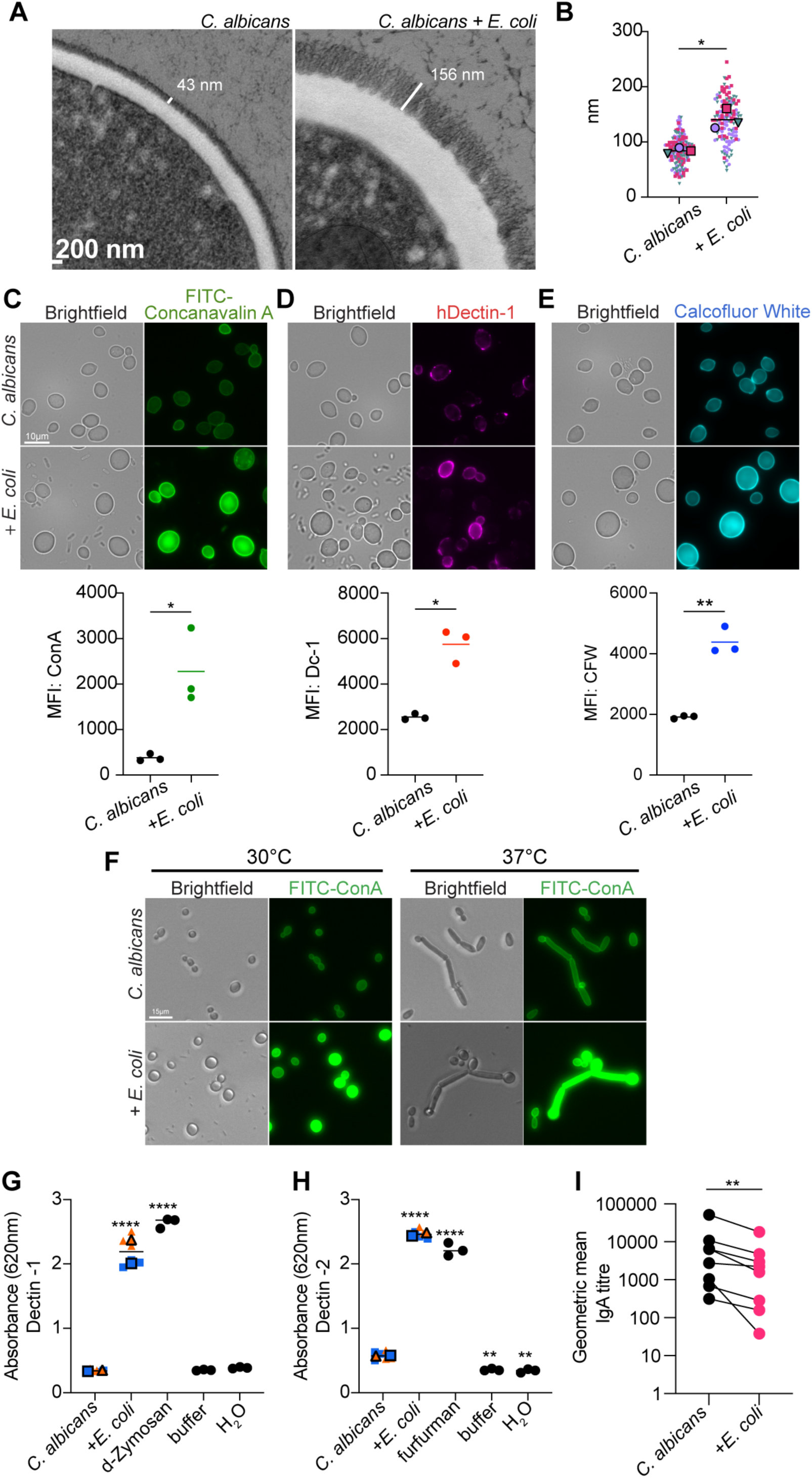
*Candida albicans* undergoes cell wall remodeling during co-culture with *Escherichia coli* with an impact on immune recognition. **A)** Representative transmission electron micrographs (TEMs) of *C. albicans* grown for 24 hours in monoculture (left) or co-culture with *E. coli* (right). White bars mark the outer mannan layer. 10,000x magnification. **B)** Quantification of mannan fibril length from TEMs across 3 biological replicates. Large outlined symbols represent the average of each biological replicate while the smaller symbols represent technical replicates. Significance determined by paired t-test on biological replicates. **C, D, E)** Determination of *C. albicans* cell wall mannan **(C)**, exposed β-1,3-glucan **(D)**, and chitin **(E)** following 24-hour co-culture with *E. coli*. Representative brightfield and fluorescent microscopy images (top). 100x magnification. Flow cytometric quantification of cell wall component (bottom). Cell wall mannan stained with FITC-Concanavalin A (ConA), exposed β-1,3-glucan stained with hDectin-1a and anti-IgG antibody conjugated with Alexa Fluor 647 (Dc-1), chitin stained with Calcofluor White (CFW). Gating strategy is illustrated in SFig 4A. Significance determined by paired t-test. MFI = mean fluorescence intensity. **(F)** Representative fluorescent microscopy images of *C. albicans* grown in monoculture or co-culture with *E. coli* at 30°C or 37°C. **G, H)** Engagement of Dectin-1 **(G)** or Dectin-2 **(H)** with *C. albicans* grown in monoculture or co-culture with *E. coli*. D-Zymosan and furfurman represent positive controls for each receptor. Large, outlined symbols represent the average of each biological replicate while the smaller symbols represent technical replicates. Significance determined by one-way ANOVA with Dunnett’s multiple comparison test. **I)** Human fecal IgA binding to *C. albicans* grown in monoculture or co-culture with *E. coli.* Significance determined by Wilcoxon test.

**Figure 2:**
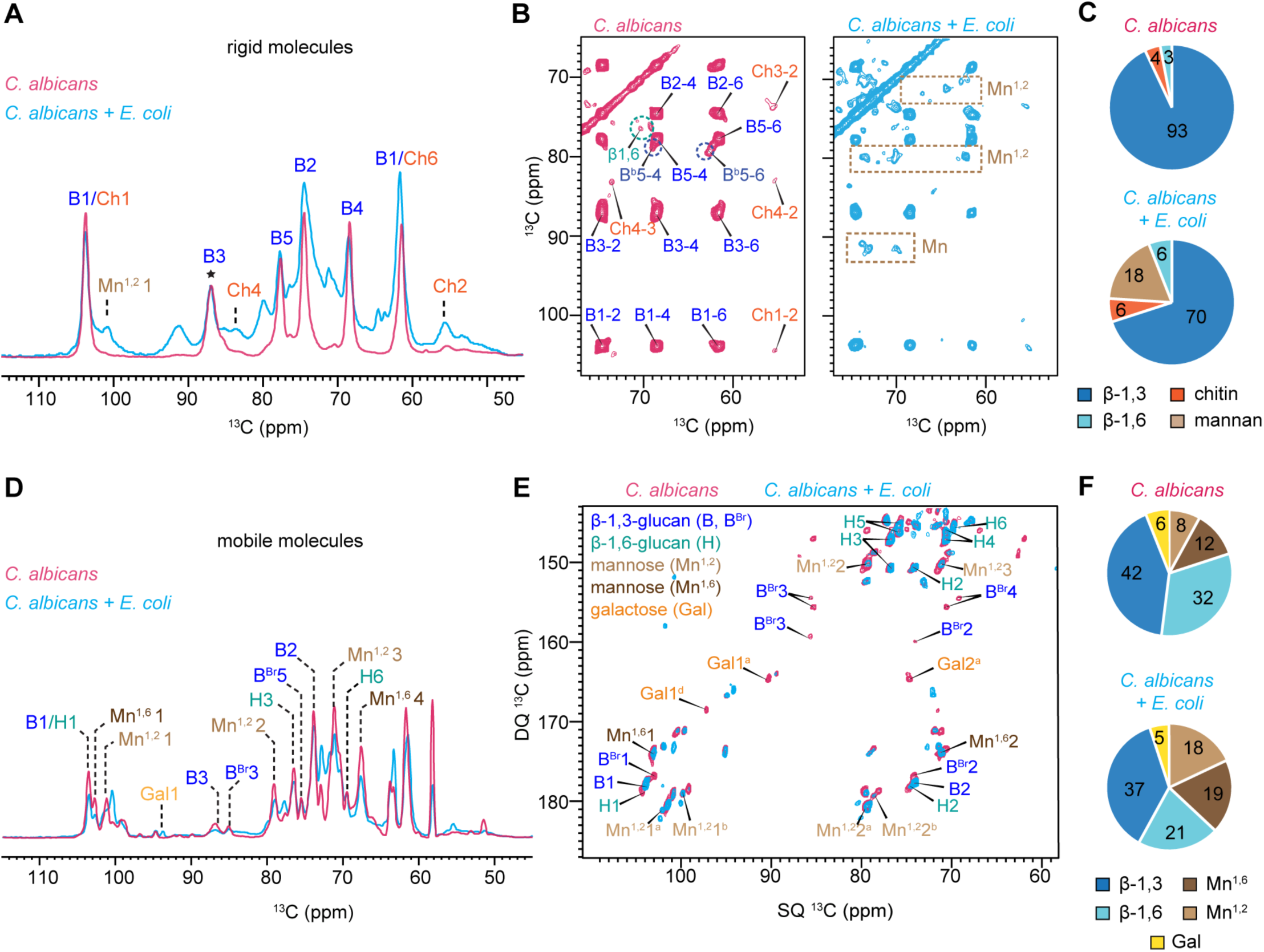
Composition of rigid and mobile polysaccharides probed by high-resolution ssNMR. **A)** Overlay of ^13^C cross-polarization (CP) spectra representing rigid glucans for *C. albicans* monoculture in pink and *C. albicans* + *E. coli* co-culture in blue. Spectra were normalized by β-1,3-glucan carbon 3 (B3; 86 ppm) peak, as indicated by the star. **B)** 2D ^13^C-^13^C CORD correlation spectrum resolving the signals of β-1,3-glucan, β-1,6-glucan, and chitin in the rigid cell walls of monoculture in pink and co-culture in blue. **C)** Molar composition of the rigid components as determined by analyzing peak volumes in 2D ^13^C-^13^C CORD spectra. Glucan types and their corresponding carbon signals are abbreviated and color-coded as follows: β-1,3-glucan (B, blue), Chitin (Ch, orange) and Mannan (Mn, light brown). **D)** Overlay of two direct polarization (DP) spectra measured with 2 s recycle delays representing mobile glucans for *C. albicans* monoculture in pink and *C. albicans* + *E. coli* co-culture in blue. **E)** Overlay of two 2D ^13^C refocused J-INADEQUATE spectra of *C. albicans* monoculture in pink and co-culture in blue, with assignments linking monomers to spectral peaks. **F)** Molar composition of these mobile components was determined by analyzing peak volumes in 2D ^13^C refocused J-INADEQUATE spectra.

After establishing our co-culture protocol, we turned to Transmission Electron Microscopy (TEM) to visualize changes in the overall cell wall architecture following co-culture with *E. coli.* Using this approach, we observed that mannan fibrils were significantly longer when *C. albicans* was grown with *E. coli* (Fig 1A-B).

TEM preparation did not consistently preserve the integrity of the inner cell wall layer, precluding our ability to measure its thickness (SFig 3). Therefore, we used cell wall staining and both fluorescent microscopy and flow cytometry to visualize and quantify components of both the inner and outer cell wall. We observed increased cell wall mannan through increased FITC-Concanavalin A (ConA) staining, increased ý-1,3-glucan exposure through increased hDectin-1a binding, and increased chitin through increased Calcofluor White (CFW) staining (Fig1C-E). Notably, during co-culture, we also observed an increase in cell size (SFig 4B), as well as heterogeneity in β-1,3-glucan exposure (SFig 5). However, we did not observe a relationship between cell size and mannan or β-1,3-glucan exposure (SFig 5A-B), although there was a positive relationship between cell size and cell wall chitin content in both monocultured and co-cultured cells (SFig 5C). In contrast, monocultured *C. albicans* cells exhibited increased chitin and β-1,3-glucan exposure primarily at bud scars, which has been previously reported ^38–40^.

**Figure 3:**
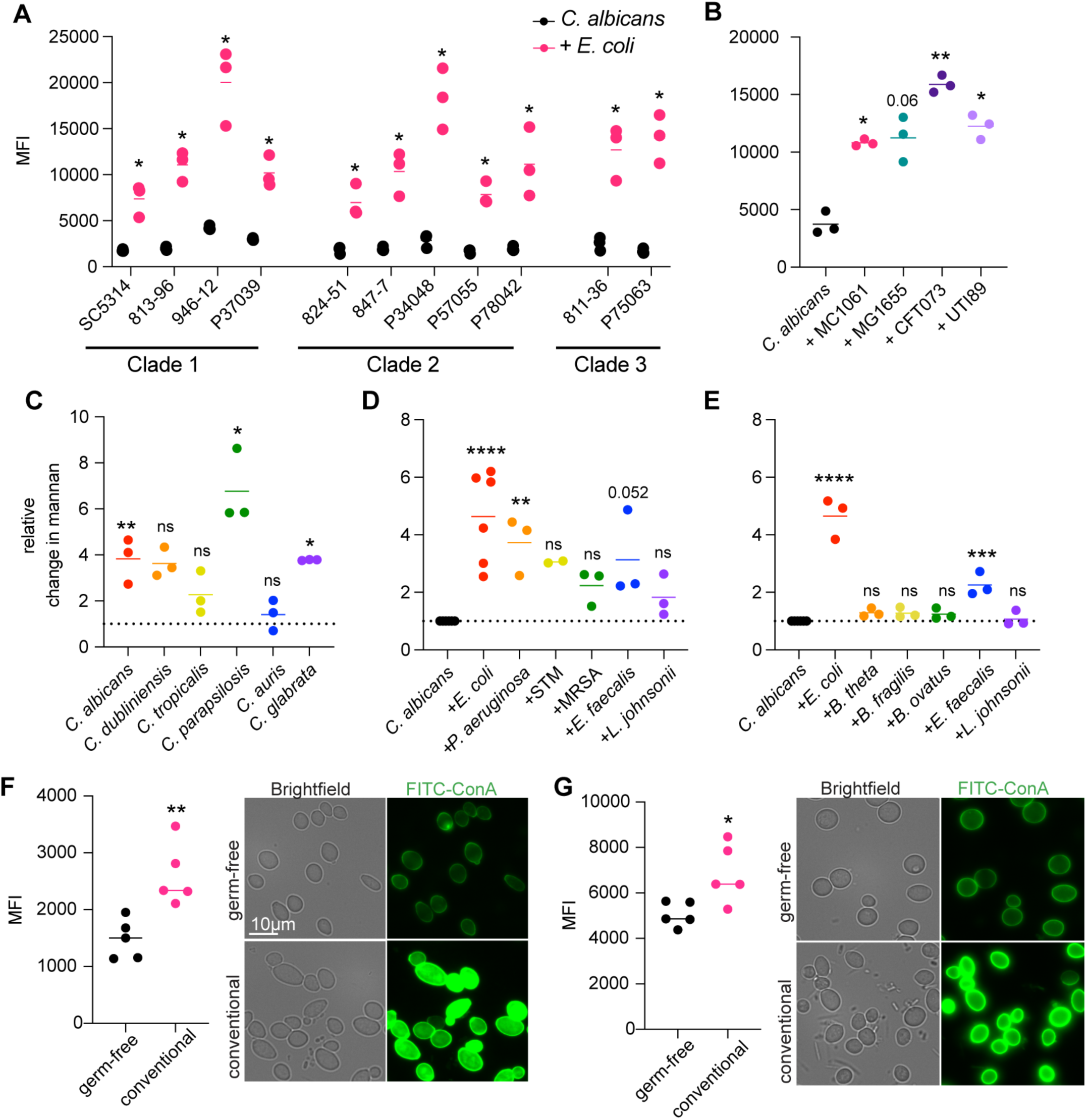
Cell wall remodeling in response to bacteria is broadly conserved. **A)** Flow cytometric quantification of cell wall mannan content of the indicated *C. albicans* isolates in monoculture or co-culture with *E. coli* after 24 hours. Significance determined by paired t-tests of each isolate during monoculture vs co-culture. **B)** Flow cytometric quantification of cell wall mannan content of *C. albicans* SC5314 following 24 hours of co-culture with the indicated *E. coli* strains. Significance determined by one-way ANOVA with Dunnett’s multiple comparison test. **C)** Relative change in cell wall mannan content of the indicated *Candida* species after 24 hours of co-culture with *E. coli.* Significance determined by paired t-tests of each *Candida* species during monoculture or co-culture with *E. coli*. **D, E)** Relative change in cell wall mannan content of *C. albicans* following 24 hours of aerobic **(D)** or anaerobic **(E)** bacterial co-culture. Significance determined by one-way ANOVA with Dunnett’s multiple comparison test. **F, G)** Flow cytometric quantification (left) and fluorescent microscopy (right) of cell wall mannan content after 24 hours of growth in germ-free or conventional mouse feces, at 37°C under aerobic **(F)** or anaerobic **(G)** conditions. Significance determined by unpaired t-test.

**Figure 4:**
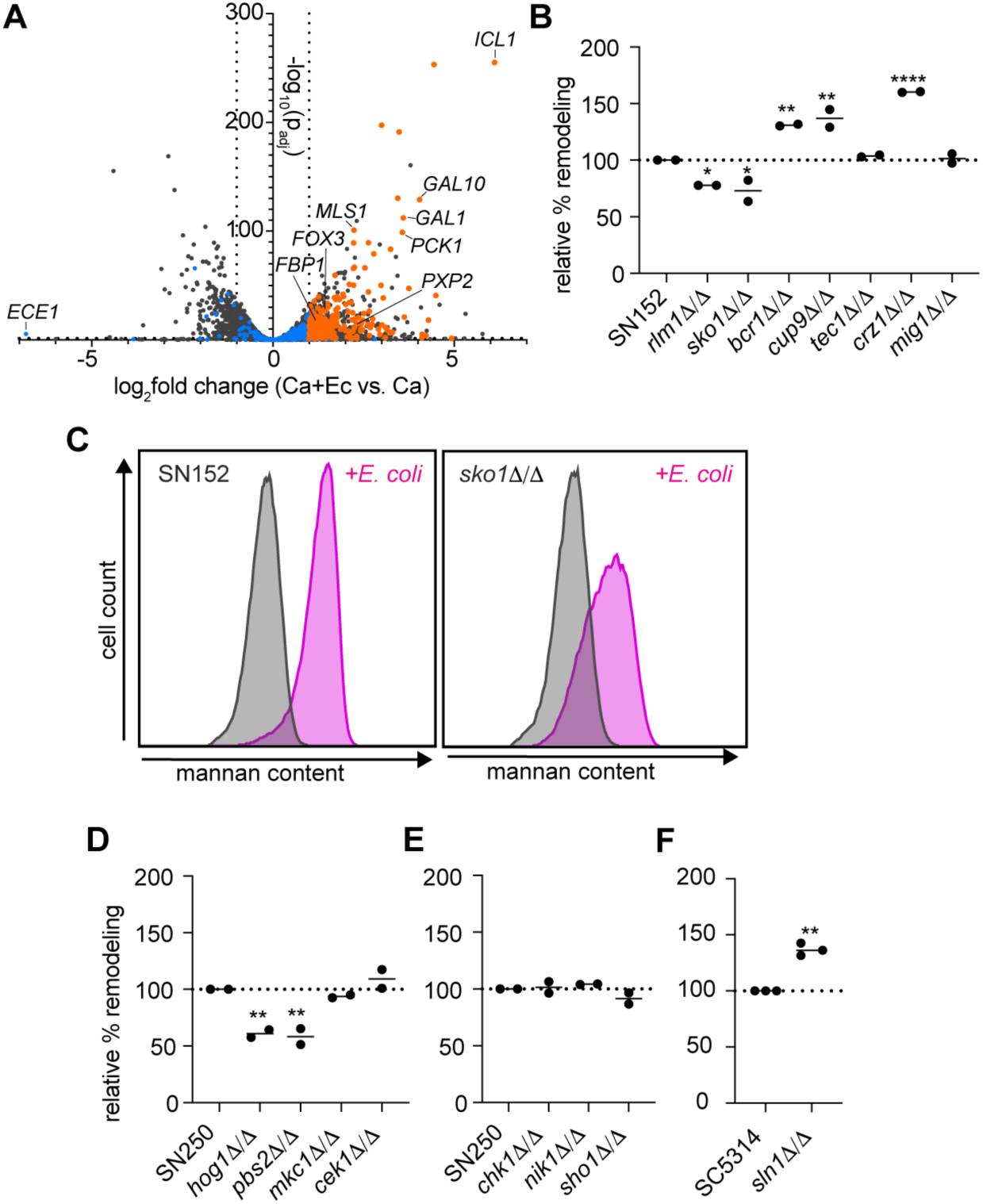
Mutants in the high osmolarity glycerol cascade are defective at remodeling in response to *E. coli* co-cure. A) Volcano plot of *C. albicans* differentially expressed genes during co-culture with *E. coli* compared to monoculture 6 hours after the initiation of co-culture. Dotted lines represent significance thresholds. Colored points are genes annotated as ’carbohydrate transport’ by g:Profiler. Orange plots are significantly upregulated and blue dots are not. **B)** Relative amount of remodeling, based on change in cell wall mannan content, following 24 hours of co-culture with *E. coli* for each indicated mutant strain from the Homann collection^63^. Set relative to cell wall remodeling of parental strain, SN152^84^. Each point is an average of two biological replicates of one independent deletion clone. Significance determined by one-way ANOVA wit’ Dunnett’s multiple comparison test. **C)** Flow cytometric histograms representing cell wall mannan content of SN152 (left) or *sko1*Δ/Δ (right) during monoculture (grey) or *E. coli* co-culture (pink). **D, E)** Relative amount of remodeling, based on change in cell wall mannan content, following 24 hours of co-culture with *E. coli* for each indicated mutant strain from the Noble collection^64^. Set relative to cell wall remodeling of parental strain, SN250. Each point is an average of two biological replicates of an independent deletion clone. Significance determined by one-way ANOVA with Dunnett’s multiple comparison test. **F)** Relative amount of remodeling, based on change in cell wall mannan content, following 24 hours of co-culture with *E. coli* for *sln1*Δ/Δ. Set relative to SC5314. Significance determined by unpaired t-test.

**Figure 5:**
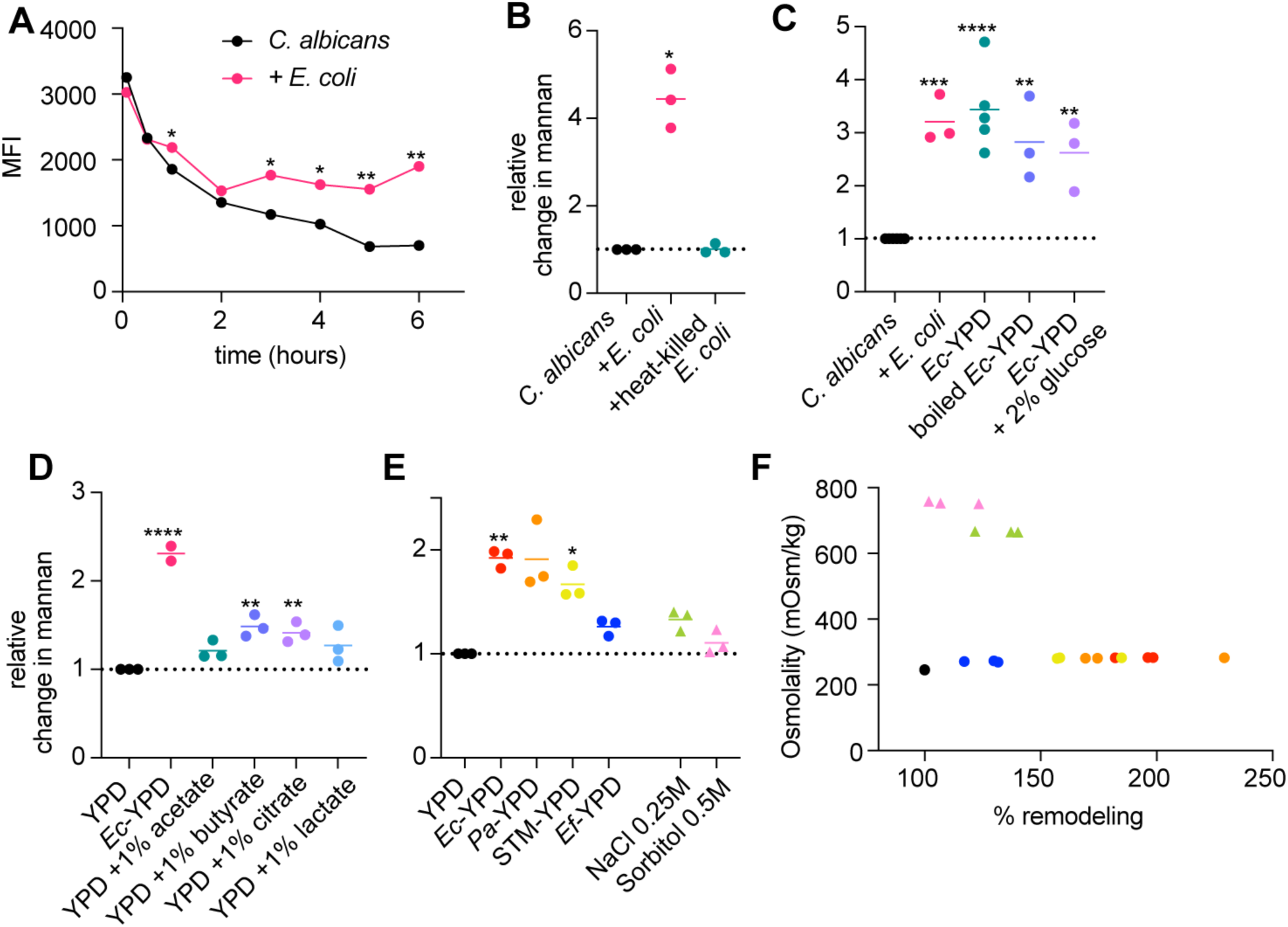
*E. coli*-induced fungal cell wall remodeling is mediated by a secreted metabolite. **A)** Flow cytometric quantification of cell wall mannan content over a 6-hour time course of *C. albicans* grown alone or in co-culture with *E. coli. C. albicans* was grown to mid-log phase before the addition of YPD media with or without *E. coli.* Significance determined by paired t-tests at each time point. **B)** Relative change in cell wall mannan content of *C. albicans* grown alone or co-cultured with live or heat-killed *E. coli* for 24 hours. Significance determined by one-way ANOVA with Dunnett’s multiple comparison test. **C)** Relative change in cell wall mannan content of *C. albicans* grown alone, with live *E. coli*, with *E. coli*-conditioned YPD, with boiled *E. coli*-conditioned YPD, or with *E. coli*-conditioned YPD with 2% glucose for 6 hours. Significance determined by one-way ANOVA with Dunnett’s multiple comparison test. **D)** Relative change in cell wall mannan content of *C. albicans* grown alone YPD, in *E. coli*-conditioned YPD, or in YPD supplemented with 1% of either acetate, butyrate, citrate, or lactate for 6 hours. Significance determined by one-way ANOVA wit’ Dunnett’s multiple comparison test. **E)** Relative change in cell wall mannan content of *C. albicans* grown alone in YPD, in conditioned YPD from indicated bacterial species, in 0.25M NaCl YPD, or in 0.5M sorbitol YPD for 6 hours. Significance determined by one-way ANOVA with Dunnett’s multiple comparison test. **F)** Linear regression comparing relative magnitude of remodeling content and osmolality of media from previous panel.

In our co-culture system, we observed primarily yeast form *C. albicans*, however, during gastrointestinal colonization in the presence of bacteria, *C. albicans* exists as a mixture of yeast and hyphal cells ^24^. Therefore, we wanted to determine if hyphal cells also undergo fungal cell wall remodeling in response to bacterial co-culture. By fluorescent microscopy, we observed that hyphal form *C. albicans* underwent cell wall remodeling in response to *E. coli* co-culture, in a similar manner as yeast cells (Fig 1F). Overall, our combination of TEM, fluorescence microscopy, and flow cytometry indicate that during co-culture with *E. coli*, *C. albicans* increases levels of cell wall mannan, chitin, and β-1-3-glucan exposure.

### *E. coli*-induced fungal cell wall remodeling impacts innate and adaptive immune responses and is relevant *in vivo*

We were particularly interested in understanding how bacterial-mediated fungal cell wall remodeling would impact innate immune recognition of *C. albicans*. Due to the observed increase of PAMPs following co-culture with *E. coli* (Fig 1A-D), we hypothesized that these cells would have more engagement with key CLRs.

Using HEK-blue reporter cells, we found that *C. albicans* cells co-cultured with *E. coli* had increased engagement with both Dectin-1 and Dectin-2 (Fig 1G-H). These results are consistent with the increase in cell wall mannan and exposure of β-1,3-glucan observed following co-culture with *E. coli* (Fig 1A-D) ^41,42^.

Beyond the clear role for the innate immune response, there is an increasing appreciation for the role of the adaptive immune responses in controlling gut commensal fungi. Intestinal IgA antibodies preferentially target *C. albicans* in the hyphal form, and this targeting promotes the competitive fitness of *C. albicans* and helps maintain homeostasis ^43^. Although we did not see an increase in hyphal formation in response to *E. coli* co-culture, we hypothesized that the altered cell wall may also impact IgA targeting. Using human fecal wash, we quantified IgA binding and found that co-culture with *E. coli* led to decreased IgA binding (Fig 1I, SFig 6). Together, this suggests that bacteria influence both innate and adaptive immune responses to *C. albicans,* with a potential impact on the inflammatory capacity of the fungus.

**Fig. 6.**
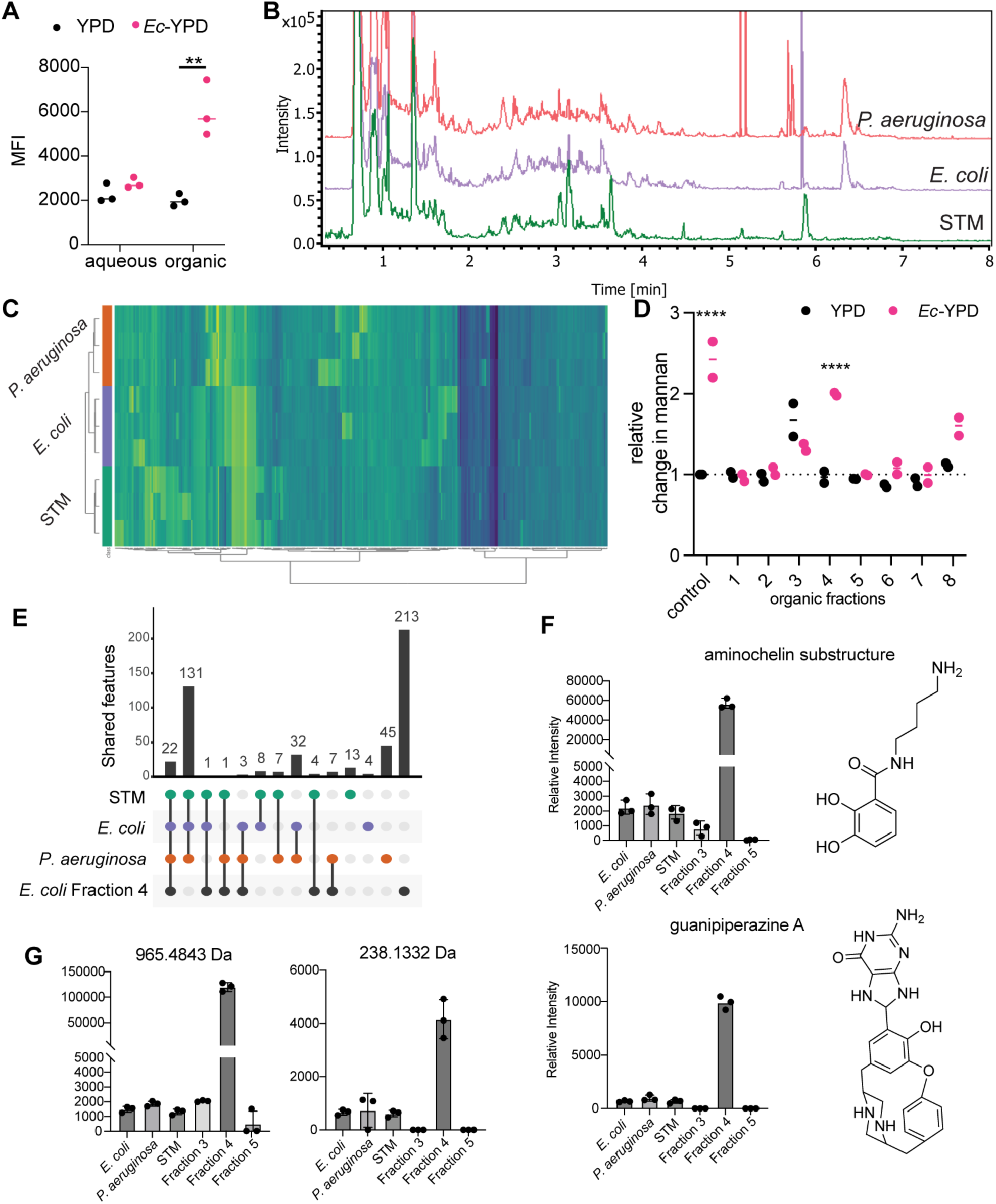
Metabolomics analysis identified secreted metabolites present in bacterial cultures that induce cell wall remodeling. A) Flow cytometric quantification of *C. albicans* cell wall mannan content after 6 hours of growth in YPD with 50 mg/mL of aqueous and organic extracts from YPD or *Ec-*YPD. Significance determined by two-way ANOVA with Sidak’s multiple comparison test. **B)** Base peak chromatograms (BPCs) of organic extracts from *P. aeruginosa*, *E. coli*, and STM as analyzed via liquid chromatography-tandem mass spectrometry (LC-MS/MS). **C)** Heatmap of LC-MS/MS data for triplicate injections of the three organic extracts of each bacterial supernatant. **D)** Relative change in cell wall mannan content for *C. albicans* grown in organic fractions of YPD or *Ec*-YPD, normalized to unfractionated YPD control. Tested concentrations of organic fractions are as follows: fraction 1, YPD 10 mg/mL, *Ec*-YPD 10 mg/mL; fraction 2, YPD 10 mg/mL, *Ec*-YPD 10 mg/mL; fraction 13 YPD 10 mg/mL, *Ec*-YPD 10 mg/mL; fraction 4, YPD 2 mg/mL, *Ec*-YPD 10 mg/mL; fraction 5, YPD 2 mg/mL, *Ec*-YPD 2 mg/mL; fraction 6, YPD 10 mg/mL, *Ec*-YPD 0.5 mg/mL; fraction 7, YPD 0.5 mg/mL, *Ec*-YPD 2 mg/mL; fraction 8, YPD 0.5 mg/mL, *Ec*-YPD 0.5 mg/mL. Significance determined by two-way ANOVA with Sidak’s multiple comparison test. **E)** Upset plot of metabolomics features shared between the three biologically active bacteria as well as the active fraction 4 from *Ec*-YPD. **F)** Two of the 22 shared features from **(E)** that followed abundance patterns that match their biological activity. These were putatively annotated as aminochelin and guanipiperazine A. **G)** Examples of unannotated shared features from **(E)** that followed the expected abundance patterns.

### Solid-State NMR reveals molecular-level remodeling of *C. albicans* cell walls during *E. coli* co-culturing

As cell wall mannan is involved in reducing exposure of β-1,3-glucan, we were somewhat surprised to see increased β-1,3-glucan exposure alongside increased mannan content ^19^. Therefore, we turned to solid-state NMR spectroscopy to characterize the molecular architecture of the *C. albicans* cell wall during monoculture or co-culture with *E. coli* ^44,45^. This method has recently been applied to a variety of pathogenic fungal species to detail their atomic-level changes in cell wall structure using living cells ^46–49^. We probed the rigid core of the fungal cell wall using 1D cross polarization (CP) and 2D ^13^C-^13^C CORD experimental schemes and this analysis revealed it was composed of β-1,3-glucan, chitin, and β-1,6-glucan, as expected (Fig 2A-B) ^49,50^. Importantly, we confirmed that there were no carbohydrate signals from *E. coli* in the rigid phase (SFig 7A). When *C. albicans* was co-cultured with *E. coli*, we observed a decrease in β-1,3-glucan content and an increase in mannan content, whereas we observed minor changes in the chitin content (Fig 2C). Interestingly, co-culture with *E. coli* shifted some mannan polymers from the mobile phase to the rigid phase, while no mannan was present in the rigid phase during monoculture (Fig 2B). We probed the mobile domain of the cell wall using ^13^C DP refocused J-INADEQUATE spectra, mostly including molecules forming the dynamic matrix and the surface layer, including β-1,3-linkages (B) in the mainchain, β-1,3,6-linkages (B^Br^) at branching points, and β-1,6-linkages (H) in the side chains (Fig. 2D-E). While we detected *E. coli* carbohydrate signals in the mobile phase, these did not overlap with the carbohydrate signals from *C. albicans* (SFig 7B). Instead, they were predominantly attributed to β-1,3-linked muramic acid, a key component of peptidoglycan (SFig 7B). In the mobile phase, we observed signals from α-1,2- and α-1,6-linked mannose residues (Mn^1,2^ and Mn^1,6^) in the mannan fibrils, as well as weak signals from galactose (Gal) residues (Fig 2D-E). Our compositional analysis revealed a decrease in β-1,3-glucan and β-1,6-glucan content and an increase in mannan content following co-culture with *E. coli* (Fig 2F). Notably, despite the decrease in overall β-glucan content, we observed an increase in exposure and immunogenicity after coculture (Fig 1D, G, H). This may relate to the finding that the β-1,3-glucan from co-culture with *E. coli* contained less branching (B^Br^) than the β-1,3-glucan from monoculture (Fig 2E). We did not observe significant changes in galactose contents between conditions (Fig 2F). This detailed characterization of the fungal cell wall provides deeper insight into how *E. coli* co-culture influences the total amount, flexibility, and branching of cell wall polysaccharides.

**Figure 7:**
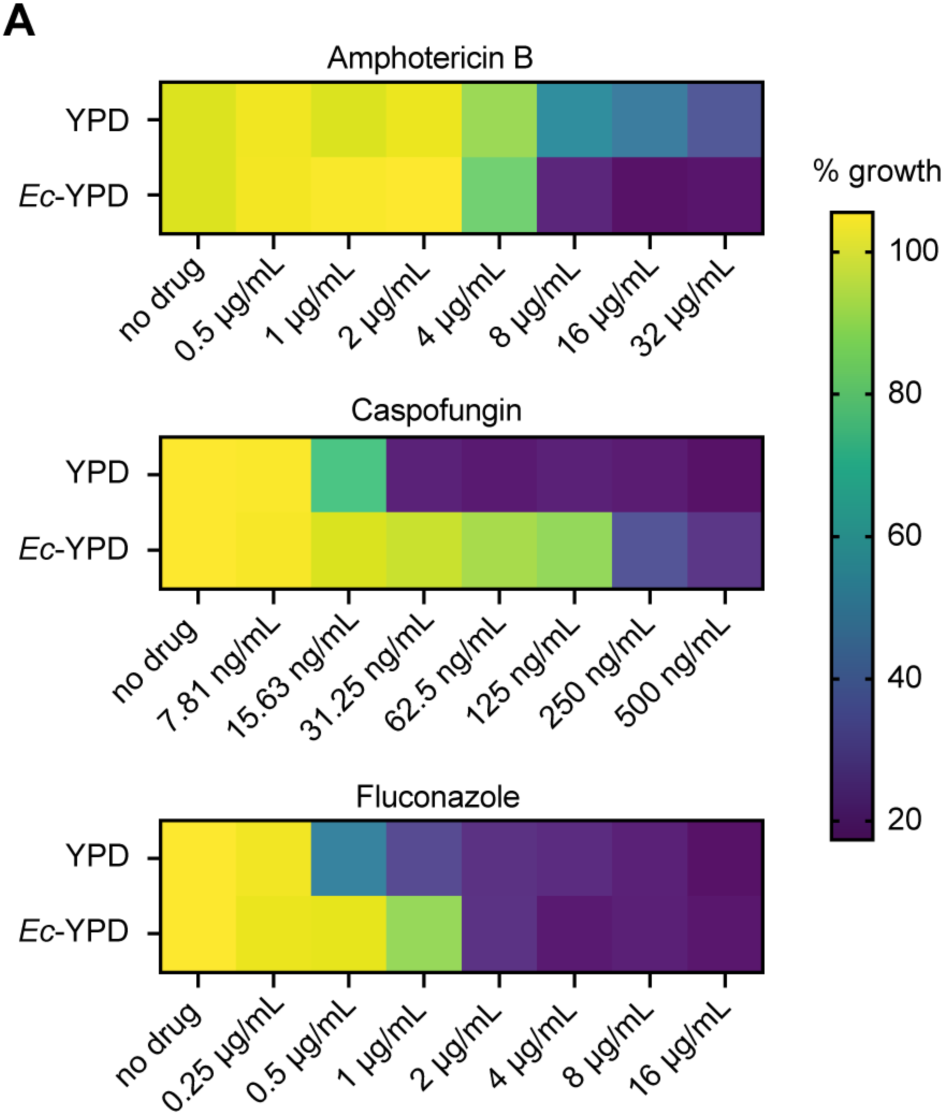
Bacterial-induced fungal cell wall remodeling alters antifungal resistance. Minimum inhibitory concentration (MIC) assay for *C. albicans* grown in YPD or *Ec*-YPD and treated with increasing concentrations of amphotericin B, caspofungin, and fluconazole.

### Fungal cell wall remodeling in response to bacteria is broadly conserved

Although the reference strain, SC5314, has been used in the majority of studies, *C. albicans* isolates display remarkable diversity in their growth rates, filamentation profiles, and interactions with host cells, among other phenotypes ^22,51–56^. Therefore, we reasoned that different genetic backgrounds of *C. albicans* may have different responses to bacterial co-culture. Using *C. albicans* isolates from commensal colonization or bloodstream infection ^22,51^, we found that regardless of genetic clade or sample origin, all tested *C. albicans* isolates underwent cell wall remodeling following co-culture with *E. coli* (Fig 3A). Additionally, we assayed two common laboratory strains and two uropathogenic isolates of *E. coli* ^57–59^ and found that all tested strains of *E. coli* induced fungal cell wall remodeling (Fig 3B). Therefore, this interaction between *C. albicans* and *E. coli* is not restricted to laboratory strains.

Next, we wanted to determine if *E. coli* induces cell wall remodeling in other *Candida* species. The tested *Candida* species had wide variation in their initial mannan content (SFig 8), and there was an overall trend towards increased cell wall mannan following co-culture with *E. coli* (Fig 3C). *C. dubliniensis* and *C. glabrata* displayed a similar magnitude of cell wall remodeling as *C. albicans*, whereas *C. tropicalis* and *C. auris* had less cell wall remodeling (Fig 3C). Meanwhile, *C. parapsilosis* displayed the greatest magnitude of cell wall remodeling following *E. coli* co-culture (Fig 3C). This indicates that *E. coli* can induce cell wall remodeling in multiple *Candida* species.

Finally, we sought to determine if fungal cell wall remodeling in response to bacteria was specific to the interaction between *C. albicans* and *E. coli* or if it could be more widely attributed to additional bacteria species. We selected a panel of bacteria that colonize similar niches as *C. albicans* and have previously reported interactions with *C. albicans* and included both gram positive and gram negative species. Under aerobic conditions, we observed a consistent trend of cell wall remodeling when *C. albicans* was co-cultured with bacteria, although the magnitude of remodeling varied between the bacterial species (Fig 3D). *E. coli* and *Pseudomonas aeruginosa* induced the most significant cell wall remodeling response, whereas the other tested species, *Staphylococcus aureus* (MRSA), *Salmonella enterica* serovar Typhimurium (STM), *Enterococcus faecalis*, and *Lactobacillus johnsonii*, induced a weaker remodeling (Fig 3D). Of the additional bacteria tested, only co-culture with *P. aeruginosa* resulted in increased cell size, similar to *E. coli* (SFig 9A).

As *C. albicans* colonizes body sites with low oxygen levels, and hypoxia is a known cue for cell wall remodeling, we also performed anaerobic co-cultures between *C. albicans* and a set of strict and facultative anaerobes ^9^. We tested the ability of three *Bacteroides* species, *B. thetaiotaomicron, B. fragilis*, and *B. ovatus* to induce fungal cell wall remodeling. *B. thetaiotaomicron* and *B. fragilis*, can use purified yeast mannan as a sole carbon source ^35^; therefore, we reasoned we might see a distinct cell wall mannan phenotype following co-culture with mannan degraders. However, co-culture with each of the three tested *Bacteroides* species, regardless of their ability to degrade fungal mannan, resulted in a consistent, but slight increase in cell wall mannan (Fig 3E). We assayed the ability of facultative anaerobes, *E. coli, E. faecalis,* and *L. johnsonii* to induce cell wall remodeling under anaerobic conditions and found that *E. coli* induced the most significant amount of remodeling, followed by *E. faecalis*, while *L. johnsonii* failed to induce remodeling (Fig 3E). We did not observe increased cell size following anaerobic co-culture with *E. coli* (SFig 9B).

As we demonstrated that multiple bacterial species can induce fungal cell wall remodeling, we wanted to extend these results to the complex community of microbes that can be found in the gut, where we hypothesize this remodeling would be happening. We obtained fecal samples from conventional and germ-free mice, resuspended them into YPD, and incubated *C. albicans* with these mixtures for 24 hours. Consistently, *C. albicans* grown in fecal samples from conventional mice had higher cell wall mannan levels than when grown in fecal samples from germ-free mice under both aerobic and anaerobic conditions (Fig 3F-G). Together, these results highlight the broad ability of physiologically-relevant bacteria to influence the fungal cell wall.

### The HOG pathway is required for fungal cell wall remodeling in response to bacterial co-culture

To determine what transcriptional changes occur during bacterial co-culture, we performed RNA sequencing on *C. albicans* grown for 6 hours in monoculture or co-culture with *E. coli*. In response to co-culture, 672 *C. albicans* transcripts were significantly upregulated and 321 were significantly downregulated (Fig 4A, Table S2). Of the upregulated transcripts, we saw a significant signature of alternate carbon utilization, indicating that *C. albicans* undergoes a global metabolic shift during co-culture with *E. coli.* Key genes involved in the glyoxylate cycle (*ICL1, MLS1*), beta-oxidation (*FOX3, PXP2*), gluconeogenesis (*PCK1*, *FBP1*), and galactose utilization (*GAL1, GAL10)*, as well as several sugar transporters and genes involved in the biosynthesis and transport of arginine were highly upregulated during co-culture. Transcripts that were significantly downregulated in response to *E. coli* largely included genes involved in DNA replication and cell cycle. This is consistent with the decreased growth rate of *C. albicans* during co-culture and potentially links cell cycle arrest with the increased cell size. Notably, we found that *ECE1*, which encodes the fungal peptide toxin candidalysin, was the most strongly downregulated gene in response to co-culture. This is in line with work from other groups demonstrating *ECE1* downregulation following treatment with supernatants from *E. coli* or *Lactobacillus* species ^60–62^.

To understand what fungal signaling pathways are involved in this cell wall remodeling response, we leveraged existing mutant libraries to select transcription factor mutants with known roles in cell wall remodeling pathways or regulation of alternate carbon metabolism ^63,64^. From our mutant analysis, we found that *bcr1*Δ/Δ, *cup9*Δ/Δ, and *crz1*Δ/Δ were hyper-responsive to co-culture with *E. coli*, indicating that they are repressors of this response (Fig 4B). However, *rlm1*Δ/Δ and *sko1*Δ/Δ had a reduced capacity to remodel during co-culture, indicating that they are positive regulators of bacterial-mediated cell wall remodeling (Fig 4B). The reduced capacity of *sko1*Δ/Δ to remodel was also apparent based on its mannan histograms, which had more overlap between conditions than that of the SN152 wildtype (Fig 4C). After identifying transcription factors required for the full remodeling phenotype, we wanted to determine which signaling pathways were responsible for transcription factor activation. Many of the relevant pathways are controlled by mitogen-activated protein kinase (MAPK) cascades; therefore, we chose representative MAPKs from three signaling pathways with well-established roles in the regulation of cell wall remodeling. We found that *mkc1*Δ/Δ, from the PKC pathway, and *cek1*Δ/Δ, from the Cek1-mediated pathway, had a normal remodeling response (Fig 4D). Representative kinases from the high osmolarity glycerol (HOG) pathway, *hog1*Δ/Δ and *pbs2*Δ/Δ, had a reduced capacity to remodel in response to *E. coli* co-culture, similar to *sko1*Δ/Δ and *rlm1*Δ/Δ (Fig 4D).

Therefore, the HOG cascade is required for fungal cell wall remodeling during co-culture with *E. coli*. This is in line with our transcription factor data, as Sko1 is associated with the HOG pathway. Finally, we sought to identify the upstream sensor responsible for activating the HOG cascade. From the existing mutant collections, we tested the involvement of two histidine kinases, Nik1 and Chk1, that feed into the HOG cascade and Sho1, an adaptor protein with a minor role in activating *C. albicans* Hog1 in response to osmotic and oxidative stress ^65^. We found that *nik1*Δ/Δ, *chk1*Δ/Δ, and *sho1*Δ/Δ had a normal remodeling phenotype, indicating that they are not responsible for activating the HOG cascade in response to *E. coli* co-culture (Fig 4E). We constructed a *sln1*Δ/Δ mutant in the SC5314 background to test the involvement of this third histidine kinase and found that *sln1*Δ/Δ was hyper-responsive to co-culture with *E. coli* (Fig 4F). This is consistent with the role of Sln1 as a negative regulator of Hog1^66,67^ and indicates that Sln1 regulates Hog1 in response to bacterial co-culture.

### Bacterial-induced fungal cell wall remodeling is mediated by secreted metabolite(s)

In addition to defining the fungal signaling pathways involved in bacterial-mediated cell wall remodeling, we wanted to understand the mechanism by which *E. coli* induces this unique phenotype. First, we measured *C. albicans* mannan content for the first 6 hours of co-culture to understand the temporal dynamics of remodeling. Fungal cell wall remodeling occurred rapidly, and the mannan content differed significantly between the conditions by 3 hours (Fig 5A). Next, we heat-killed *E. coli* prior to the initiation of co-cultures and we did not observe cell wall remodeling under these conditions, indicating that *E. coli* must be live and metabolically active to induce fungal cell wall remodeling (Fig 5B). To determine if a secreted factor is responsible for the induction of fungal cell wall remodeling, we grew *E. coli* overnight in YPD before bacterial cells were removed by filtration to create *E. coli*-conditioned YPD (*Ec*-YPD). We found that 6 hours of growth in *Ec*-YPD was sufficient to induce cell wall remodeling to the same degree as co-culture with live *E. coli* (Fig 5C). This phenotype remained when *Ec*-YPD was heated to 60°C for 30 minutes to inactivate proteins, indicating that a secreted, heat stable factor mediates the induction of cell wall remodeling (Fig 5C). We observed a reduced growth rate when *C. albicans* was grown in *Ec*-YPD, consistent with results from live co-culture and RNAseq data (SFig 10). We also considered that nutrient deprivation could be triggering fungal cell wall remodeling and therefore supplemented Ec-YPD with 2% glucose, to ensure *C. albicans* had sufficient access to its preferred carbon source. The cell wall remodeling response was conserved following growth in *Ec*-YPD + 2% glucose, indicating that glucose competition is not responsible for the cell wall remodeling phenotype (Fig 5C).

As bacteria secrete metabolic byproducts, including short chain fatty acids (SCFAs), that could influence the *C. albicans* cell wall, and as our RNAseq experiment identified a global metabolic remodeling signature during *E. coli* co-culture, we considered that changes in the available carbon sources could be involved in the remodeling phenotype ^12,13,28^. We added 1% w/v of acetate, butyrate, citrate, or lactate to YPD and compared cell wall mannan content following growth in these conditions to *Ec-*YPD. None of the added organic acids induced fungal cell wall remodeling to the same magnitude as *Ec-*YPD (Fig 5D). Therefore, the organic acids tested here are not drivers of bacterial-mediated cell wall remodeling.

Due to the involvement of the HOG pathway in the response to fungal cell wall remodeling (Fig 4D), we wanted to determine if our remodeling phenotype was driven by osmotic stress. Exposure to 1M NaCl has previously been shown to induce short-term changes in cell wall architecture, but the impact of prolonged growth under osmotic stress on the cell wall has not been described ^14^. To examine the relationship between osmolality and remodeling, we compared the response to either two known osmotic stressors, NaCl and sorbitol, pr conditioned media from those bacteria that we previously demonstrated to induce remodeling (Fig 5E). Each of these cues was able to induce remodeling, albeit to varying degrees (Fig 5E). We then measured the osmolality of each of these media conditions but found no relationship between osmolality and extent of cell wall remodeling, indicating that despite the involvement of the HOG cascade, osmotic stress is not driving bacterial-mediated cell wall remodeling (Fig 5F).

This led us to hypothesize that a potentially uncharacterized secreted bacterial metabolite was responsible for inducing *C. albicans* cell wall remodeling. To start to characterize the secreted metabolome, we turned to bioassay-guided fractionation of *Ec*-YPD, starting with differentiating between aqueous and organic fractions. Only the organic extracts were able to induce cell wall remodeling when added to YPD (Fig 6A). Since conditioned media from *E. coli, P. aeruginosa*, and STM were all able to induce remodeling (Fig 5E), we compared the organic extracts from each organism using untargeted metabolomics via liquid chromatography-tandem mass spectrometry (LC-MS/MS). Based on a comparison of the base peak chromatograms (Fig 6B), and heat map of the metabolome (Fig 6C), we identified shared metabolomic features between all species. While some features remain well conserved and highly abundant in all samples, other features were present in varying amounts. Given the large number of shared metabolites, we then further fractionated the organic extract from *Ec*-YPD and tested the ability of each fraction to induce cell wall remodeling compared with fractions of the media control. We observed that *Ec*-YPD fraction 4 induced the most significant cell wall remodeling (Fig 6D). Comparing the features between the bacterial extracts and *Ec*-YPD fraction 4, there were only 22 shared features (Fig 6E), which we prioritized for further identification. We then compared metabolite abundance patterns (SFig 11) with the biological activity, looking specifically for metabolites with similar abundances in the bacterial strain extracts with high abundance in *Ec*-YPD fraction 4 and lower abundances in *Ec*-YPD fractions 3 and 5. Nine of the 22 prioritized metabolites followed this abundance pattern, including two metabolites putatively annotated as an aminochelin substructure and guanipiperazine A (Fig 6F), as well as unannotated metabolites (Fig 6G, SFig 11).

### Bacterial-mediated fungal cell wall remodeling modulates antifungal resistance

As most available classes of antifungals function through targeting cell wall or cell membrane components, we sought to determine if bacterial-induced cell wall remodeling could modulate antifungal drug susceptibility. We determined the minimum inhibitory concentration (MICs) for amphotericin B, caspofungin, and fluconazole, in both YPD and *Ec-*YPD. Growth in *Ec-*YPD led to a subtle, but reproducible increase in sensitivity to amphotericin and a slight increase in resistance to fluconazole (Fig 7A, SFig 12). Most strikingly, we found a significant increase in resistance to caspofungin when *C. albicans* was grown in *Ec-*YPD (Fig 7, SFig 12). The observed increased resistance to caspofungin is in agreement with work showing that increased cell wall chitin levels correlate with increased echinocandin resistance ^68,69^. This highlights how interactions with bacteria may alter the efficacy of antifungal treatment.

## Discussion

The fungal cell wall is a dynamic and essential organelle that provides structure, protects the cell from external stresses, and mediates interactions with host immune cells. As a result of its important roles, the fungal cell wall is highly responsive to changes in environmental conditions. Here, we have demonstrated that *C. albicans* undergoes cell wall remodeling following co-culture with *E. coli* that increases cell wall mannan and chitin content as well as unmasking the major immunostimulatory ligand, β-1,3-glucan. Solid-state NMR analysis of the rigid and mobile phases of the fungal cell wall revealed changes in cell wall content, polymer flexibility, and branching. These significant changes in the cell wall composition resulted in altered antifungal susceptibility, increased recognition by innate PRRs, and decreased binding of human IgA. We found that fungal cell wall remodeling is broadly conserved amongst *Candida* and bacterial species, and we demonstrated that communities of microbes derived from conventional mice fecal samples induce fungal cell wall remodeling, highlighting the relevancy of this phenotype in gastrointestinal colonization. Through mutant analysis we identified that the HOG cascade in *C. albicans* is required for the full remodeling response and demonstrated that *C. albicans* undergoes a global metabolic shift during co-culture with *E. coli*. Finally, we determined that bacterial-mediated fungal cell wall remodeling is mediated by secreted metabolite(s) and used untargeted metabolomics coupled with bioassay-guided fractionation to explore the metabolome from species that are able to induce remodeling. Overall, our work reveals that bacteria influence the *C. albicans* cell wall with implication for immune recognition and antifungal resistance.

This work adds to existing knowledge of environmental cues that trigger cell wall remodeling in *C. albicans* and alter exposure of the major fungal PAMP, β-1,3-glucan ^9–15^. A potential limitation of our work was that we primarily focused on growth under rich media conditions and in the absence of other host-associated cues that are known to influence fungal morphology and cell wall composition. In the future, it will be important to characterize how the presence of multiple cell wall remodeling cues, including the impact of different bacterial compositions, impacts the fungal cell wall and how these cues are prioritized. For example, hypoxia, which triggers β-glucan masking, was shown to be the dominant cue over exposure to CO_2_, a cue that induces unmasking of β-glucan ^70^. Understanding how combinations of known cell wall remodeling cues impact cell wall composition and other important aspects of fungal biology and how response to these cues differ *in vitro* and *in vivo* will be important topics of future research, as *C. albicans* is exposed to many of these cues in parallel during host colonization. In line with this idea, there is a growing appreciation that host-associated cues have profound impacts on fungal biology and that these effects are often not accounted for during *in vitro* research. Recent work demonstrated that host-relevant concentrations of CO_2_ alters resistance to echinocandins, emphasizing that antifungal susceptibility testing in a laboratory environment can preclude identification of antifungal resistance ^71^. Studies have also demonstrated that the presence or absence of gastrointestinal bacteria in murine models can influence fungal evolution, colonization, and virulence ^5,23,29,72,73^. Overall, these results highlight the importance of conducting research in conditions that most closely mimic relevant *in vivo* conditions for the greatest interpretability of data in relation to human colonization and disease.

Curiously, our cell wall staining and flow cytometric analysis of the cell wall revealed that mannan content and β-1,3-glucan exposure were increased during *E. coli* co-culture. This observation is at odds with the model that cell wall mannan masks β-1,3-glucan from detection by immune cells ^19^. When we analyzed the cell wall in more depth through SS-NMR, we discovered that *E. coli* co-culture led to an increase in cell wall mannan but a reduction in β-glucan. Additionally, co-culture with *E. coli* changes the physical properties of these polysaccharides. Some mannan polymers from *E. coli* co-culture had shifted from the mobile phase to the rigid phase, and β-glucan from *E. coli* co-culture was more linear. Therefore, the relationship between the total amount of cell wall polysaccharides and exposure of β-1,3-glucan is not necessarily straightforward. Our detailed analysis of the fungal cell wall using multiple, parallel methods underscores the complexity of this organelle and suggests that properties including polymer length and branching, flexibility, porosity are all important determinants of how the cell wall is recognized by immune cells.

We identified that the transcription factors, Rlm1 and Sko1, as well as the central kinases of the HOG cascade, Hog1 and Pbs2, were required for the full remodeling phenotype. Deletion of these regulators resulted in a 25-40% decrease in the magnitude of remodeling but did not abolish the phenotype. Cell wall remodeling is complexly regulated and frequently involves crosstalk and coordination between multiple signaling pathways and the molecular response to different environmental stimuli is unique ^74–76^. Therefore, it is highly probable that additional signaling pathways are involved in regulating bacterial-mediated fungal cell wall remodeling. Although we identified that the histidine kinase, Sln1, is involved in initiating the signaling cascade, the HOG cascade could also be regulated through crosstalk with additional pathways.

We also identified several metabolites with abundance patterns consistent with their parent sample’s capacity for fungal cell wall remodeling. While most of these metabolites remain unknown, three were putatively annotated as an aminochelin substructure, guanipiperazine A, and asperorydine N. Aminochelin is a catecholamine siderophore originally isolated from *Azotobacer vinelandii*^77^ grown in iron-limited conditions and is a substructure of several larger siderophores isolated from the same strain^78^. While the role of these intriguing siderophore molecules in the interaction between *E. coli* and *C. albicans* has not yet been confirmed, it is intriguing to consider what role these small molecule siderophores might play in fungal cell wall remodeling. In data not shown, iron supplementation was not sufficient to reverse the remodeling, suggesting it is not merely a lack of iron that leads to remodeling. Guanipiperazine A was originally isolated via genome mining from *Streptomyces chrestomyceticus*^79^ and was subsequently shown to have antifungal activity^80,81^, although that mechanism is undefined. Interestingly, asperorydine N is part of a series of metabolites originally isolated from *Aspergillus* strains, including *A. flavus* and *A. oryzae*^82,83^. Given that we have putatively identified this molecule from our bacterial strains, it is possible that the bacteria are metabolizing compounds that were initially present in the yeast extract that they were grown in. Due to the complex mixture of many unknown metabolites, future work will be needed to examine and further isolate all prioritized metabolites to determine their role(s) in fungal cell wall remodeling.

## Materials and Methods

### Strains and culture conditions

The fungal and bacterial strains used in this study are listed in Supplemental Table S1. *C. albicans* strains were grown in liquid YPD (1% yeast extract, 2% bacto-peptone, 2% glucose) media at 30°C. *E. coli*, *P. aeruginosa*, and STM were grown in liquid LB media at 37°C. MRSA was grown in liquid TSB media at 37°C. *E. faecalis* and *L. johnsonii* were grown in liquid MRS media at 37°C. *Bacteroides* species were grown in TYG media at 37°C in an anaerobic chamber.

### Fungal-bacterial co-culture

Overnight cultures of *C. albicans* were diluted in YPD to an OD_600_ of 0.1. Overnight cultures of bacteria were washed with water to remove bacterial media, then diluted to an OD_600_ of 0.5 in YPD. 1 mL of the *C. albicans* dilution was added to each well of a 6- or 12-well plate. 1 mL of fresh YPD or 1 mL of the bacterial dilution was added to the *C. albicans* dilution. Co-cultures were conducted at 30°C in aerobic conditions or at 37°C in an anaerobic chamber. For hyphal co-cultures, *C. albicans* was grown aerobically at 37°C for 3 hours before the addition of fresh YPD or *E. coli*. Cells were collected after 24 hours, unless otherwise indicated. The cell suspensions were collected in 1.5 mL microcentrifuge tubes and centrifuged at 13,000 rpm for 1 minute to pellet cells. The supernatant was removed, and cells were fixed with 1 mL of 4% paraformaldehyde (PFA) for 10 minutes at room temperature. The cells were centrifuged again and PFA was removed. The cells were washed with 1 mL of PBS 3 times. Cells were resuspended in 1 mL of PBS and stored at -20°C.

### Transmission electron microscopy

Co-cultures were performed as described above, and fresh cells (not fixed) were provided to the Michigan Microscopy Core for preparation. Samples were mixed 1:1 with YPD media and 1% ultra low gelling temperature agarose for a final concentration of 0.5% agarose. These suspensions were frozen with a Lecia EM ICE high pressure freezer and stored in liquid nitrogen. Frozen samples were removed from liquid nitrogen and freeze substituted in a solution containing 2% OsO_4_, 0.1% uranyl acetate, and 5% H_2_O in a Lecia AFS-2. Freeze substitution was carried out at the following temperature schedule: keep samples at - 90°C for 24 hours, raise temperature to -60°C in 15 hours (2°C/hour), keep samples at -60°C for 24 hours, raise temperature to -30°C in 15 hours (2°C/hour), keep samples at -30°C for 24 hours, raise temperature to 4°C at 3°C/hour, raise temperature to 25°C in 30 minutes, keep samples at 25°C for 1 hour. After freeze substitution, samples were washed with acetone 3 times then infiltrated with Epon resin/acetone gradient solutions at the following schedule: samples infiltrated with 1:5 solution for 3 hours, 1:2 solution for 4 hours, 1:1 solution overnight, 2:1 solution for 8 hours, 5:1 solution overnight, and pure resin for 24 hours. Samples were embedded in fresh resin and cured at 60°C for 48 hours. Cured samples were cut into 70 nm sections and mounted on Formvar/carbon copper grids for imaging. Samples were imaged using a JEOL JEM 1400 Plus Transmission Electron Microscope. ImageJ software was used to determine the length of mannan fibrils. For each image, the scale was set using the embedded scale bar. For each cell, 3 measurements were taken for mannan fibril length and these measurements were averaged for final fibril length. A minimum of 35 cells were measured for each replicate.

### Flow cytometric quantification of cell wall components

Cells from co-culture experiments were thawed and transferred to a new microcentrifuge tube for staining. To quantify mannan content, cells were stained with 5 μg/mL of FITC-Concanavalin A (MilliporeSigma, C7642) for 30 minutes. To quantify exposed β-1,3-glucan, cells were blocked with 3% bovine serum albumin & 5% normal goat serum (Invitrogen, 10000C) for 30 minutes. After blocking, cells were stained with 15 ug/mL of hDectin-1a (InvivoGen, fc-hec1a-2) for 1 hour. Cells were washed twice with PBS before secondary staining with 4 mg/mL of goat raised anti-human IgG antibody conjugated with Alexa Fluor 647 (Invitrogen A-21445) for 30 minutes. To quantify chitin content, cells were stained with 0.1 g/L of Calcofluor White (MilliporeSigma, 18909-100ML-F) for 10 minutes. Following staining, cells were washed with 500uL PBS three times and resuspended in 500uL PBS. Samples were analyzed on a LSRFortessa Flow Cytometer (BD Bioscience, NJ, USA) using BD FACSDiva Software. 100,000 events were recorded for each sample. FlowJo software was used to gate for fungal cells as shown in SFig 4A and to determine the mean fluorescence intensity (MFI).

### Fluorescence Microscopy

Cells were stained as described above. Microscopy was performed on a Biotek Lionheart FX automated microscope using 100X oil objective or 40X air objective. The FITC imaging filter cube was used to visualize mannan, the DAPI imaging filter cube was used to visualize chitin, and the CY5 imaging filter cube was used to visualize exposed β-1,3-glucan.

### HEK Cells Methods

HEK-Blue mDectin-1b, mDectin-2, and control Null1-v cells were purchased from Invivogen and cultured in DMEM with 100 U/mL Pen/Strep, 2 mM L-glut, 10% heat-inactivated fetal bovine serum, and 100 µg/mL Normocin. The following selective antibiotics were used for each cell line: mDectin-1b, 1 µg/mL puromycin, 1x HEK-Blue CLR Selection; mDectin-2, 1 µg/mL puromycin, 1x HEK-Blue CLR Selection, 30 µg/mL blasticidin; Null1-v, 100 µg/mL zeocin. For in vitro reporter cell assays, formaldehyde fixed *C. albicans* strain SC5314 alone, co-cultured with *E. coli*, and *E. coli* alone were added to 5.04×10^4^ HEK-Blue cells at an MOI of 2 (based on *C. albicans*) in 96-well plates in HEK-Blue Detection medium. Control agonists were included for each cell line. Depleted zymosan (100 µg/mL) was used as a positive control for mDectin-1b, furfurman (1000 µg/mL) was used as a positive control for mDectin-2, and trehalose-6,6-dibehenate (TDB) (100 µg/mL) was employed as a negative control for both cell lines. All three control agonists were plated with the Null1-v cells. The cultures were incubated for 20 hours at 37°C in 5% CO2, followed by quantification of secreted embryonic alkaline phosphatase (SEAP) by measuring absorbance at 620 nm.

### Quantification of IgA binding

To quantify IgA binding to monocultured or co-cultured *C. albicans*, 10 µl of 10^7^ *C. albicans/*ml were incubated for 20 min at 4°C with 25 µl cleared fecal homogenate from healthy human donors^43^ diluted 1:4 in PBS + 1% BSA. Cells were centrifuged for 5min at 3000rpm and washed 2 times with 150ul PBS + 1% BSA. Cells were then stained for 15 min with 50 ul anti-human IgA (Jackson ImmunoResearch 109-605011) at a 1:500 dilution. Cells were centrifuged for 5min at 3000 rpm, and washed 2 times with 150ul PBS + 1% BSA, and analyzed by flowcytometry using the CYTOflex flow cytometer. Geometric mean binding intensity of IgA was normalized to both monoculture or co-cultured *C. albicans* stained with anti-human IgA only (geometric mean IgA binding of fecal-wash-stained minus no fecal wash control).

### Solid-State NMR

1D and 2D high-resolution solid-state NMR experiments were conducted on a Bruker Advance NEO 800 MHz (18.8 Tesla) spectrometer at MSU Max T. Roger NMR facility. The experiments were conducted using a 3.2 mm triple-resonance HCN probe under 15 kHz magic-angle spinning (MAS) at 280 K. The ^13^C chemical shifts were externally referred to adamantane CH_2_ signal at 38.48 ppm on the tetramethylsilane (TMS) scale. The magic angle was calibrated using KBr. Typical ^1^H radiofrequency field strengths 50-83 kHz and 50-62.5 kHz for ^13^C. The initial magnetization for the experiment was created using ^1^H-^13^C cross-polarization that preferentially detects rigid molecules. Typically, 1 ms Hartmann-Hahn contact was used for CP. The 2D refocused J-INADEQUATE experiment ^85^ was integrated with either direct polarization (DP) and short recycle delays of 2 s for the selective detection of mobile components. A 2D ^13^C-^13^C correlation experiment was also measured using the CORD ^50^ scheme with a 53-ms recoupling period. The composition of rigid and mobile components was determined by analyzing peak volumes in 2D CORD and DP refocused J-INADEQUATE spectra, respectively. To minimize uncertainties from spectral crowding, only well-resolved signals were considered for the compositional analysis. Resonance assignment was based on the values reported for *C. albicans* and other *Candida* species^49^.

### Fecal Samples

Fecal samples were manually collected from 10 germ-free and 10 conventional mice. Germ-free samples were confirmed via PCR. One fecal pellet was resuspended in 2mL YPD and *C. albicans* was added to this mixture at an OD of 0.1. *C. albicans* was incubated for 24 hours at 37°C under aerobic or anaerobic conditions before samples were collected and analyzed as outlined above.

### RNA Extraction

RNA extraction was performed using a formamide extraction method ^86^. Briefly, cells were grown in monoculture or in co-culture with *E. coli* in YPD at 30°C for 6 hours before being harvested by centrifugation and all media was removed. Dry cell pellets were frozen on dry ice and stored at -80 °C before processing. To extract RNA, cell pellets were thawed at room temperature and resuspended in 100 μL FE Buffer (98% formamide, 0.01M EDTA). 50 μL of 500 μm RNAse-free glass beads was added to this suspension and the mixture was homogenized for 30 sec 3 times using a BioSpec Mini-Beadbeater-16 (Biospec Products Inc., Bartlesville, OK, USA). The resulting cell lysate was clarified by centrifugation to remove cell debris. The supernatant was collected as the crude RNA extract. The crude extract was purified using a Qiagen RNeasy mini kit (ref 74104, Qiagen) according to the manufacturer’s instructions. Samples were DNAse treated with Invitrogen DNAse (RNAse free) (Qiagen, cat no. 79254). The integrity and purity of the extracted RNA was confirmed via Nanodrop and agarose gel electrophoresis using the bleach gel method prior to downstream applications ^87^. Sequencing libraries were prepared and sequenced on an Illumina NovaSeq 6000 with 150 bp paired-end reads by Novogene.

### RNAseq analysis

RNA-seq analysis was performed in Galaxy (usegalaxy.org). Reads were evaluated using FastQC and trimmed using trimmomatic^88^, followed by quantification of transcripts from the SC5314 using Kallisto^89^. Differential gene expression between *C. albicans* in monoculture or *E. coli* co-culture (Table S2) was compared using DESeq2^90^. Gene ontology analysis was performed on significantly upregulated or downregulated genes using g:Profiler^91^. All data are available on NCBI SRA at PRJNA1290278.

### Construction of *sln1*Δ/Δ mutant

*SLN1* was deleted from the SC5314 strain background using a transient CRISPR approach [cite]. The *SLN1::NAT* cassette was amplified from the NAT flipper plasmid [cite] using primer pair oTO2610 & oTO2611 and Cas9 was amplified from [plasmid] using primer pair oTO40 & oTO41. The *SLN1* specific sgRNA guide was created by amplifying two segments from the [plasmid] using primer pairs oTO2613 & oTO6 and oTO2612 & oTO8 then the segments were fused using the nested primer pair oTO7 & oTO9. Integration was tested using primers pairs oTO3 & oTO2614 and oTO6 & oTO2615 and loss of the wild-type *SLN1* gene was tested using oTO2614 & oTO2730. Plasmids and oligonucleotides used in this study are listed in Tables S3 and S4.

### *E. coli*-conditioned YPD

*E. coli* was grown for 18 hours in YPD media at 37°C. Cells were harvested by centrifugation at 4000 rpm for 5 minutes for small quantities (5mL) or at 10,000xg for 10 minutes for large quantities (500mL). The supernatant was collected and passed through a 0.2 μm filter to remove any remaining cells. The solution was used immediately or aliquoted and stored at -20°C to be used within 2 weeks.

### Determination of media osmolality

Conditioned YPD media from *P. aeruginosa,* STM, and *E. faecalis* was prepared as described above. The osmolality of blank YPD media, bacteria-conditioned YPD media, and YPD media with the addition of 0.25 M NaCl or 0.5 M sorbitol were measured using a Fiske Model 210 Micro-Osmometer.

### Extraction and fractionation

Solvents used in extractions were all high-performance liquid chromatography (HPLC) grade from Sigma-Aldrich. Three rounds of sequential ethyl acetate extractions were performed on supernatants of bacterial cultures grown for 24 hours. For metabolomics analysis, 75 mL cultures of *E. coli*, *P. aeruginosa*, STM, and unconditioned YPD media were extracted. For fractionation, large scale cultures (500 mL) of unconditioned YPD media or of *E. coli* conditioned YPD media were extracted. Bacterial cultures were sonicated and extracted with equal volumes of ethyl acetate three times, each for 15 min incubation at room temperature. For each extract, the aqueous and ethyl acetate layers were separately combined, dried using rotary evaporation, transferred to vials, and stored at -80°C prior to further analysis.

Organic extracts from unconditioned YPD media and *E. coli* conditioned media (*Ec*-YPD) (144 mg and 138 mg, respectively) were separately added to Biotage Sfar C_18_ D columns connected to a Biotage Selekt for fractionation. Eight fractions were collected using a 0-100% methanol (MeOH)/water solvent gradient with 2 column volumes (18 mL) per fraction resulting in fraction 1 (0-20% MeOH), fraction 2 (20-30% MeOH), fraction 3 (30-40% MeOH), fraction 4 (40-50% MeOH), fraction 5 (50-60% MeOH), fraction 6 (60-70% MeOH), fraction 7 (70-90% MeOH), and fraction 8 (90-100% MeOH). Fractions were collected, dried using a Biotage V10, transferred to vials, and stored at -80°C prior to further analysis.

### Metabolomics data acquisition and analysis

Solvents used for LC-MS/MS were of LC-MS grade and sourced from Sigma-Aldrich. To prepare for analysis, samples were resuspended in 50% MeOH:water to a concentration of 1 mg/mL. LC-MS/MS analysis was performed using an Acquity UPLC HSS-T3 C_18_ 1.8 µm column with an Agilent 1290 Infinity II Bio UHPLC attached to a Bruker timsTOF Pro2 Bruker mass spectrometer. Each sample was injected in technical triplicate at random. Samples were eluted using a 0.3 mL/min 9.5-minute solvent gradient of 0.1% formic acid in water (mobile phase A) and 0.1% formic acid in acetonitrile (mobile phase B). The solvent gradient conditions were as follows: 0.5 min hold at 5% B, 1.5 min ramp to 20% B, 1 min ramp to 60% B, 3 min ramp to 100% B, 2 min hold at 100% B, 0.5 min ramp back to 5% B, and a re-equilibration hold at 5% B for 1 min. Data was acquired using positive ionization mode with a VIP-HESI (Vacuum Insulated Probe Heated Electrospray Ionization) source using the following conditions: collision energy of 10eV, capillary voltage of 4500 V, dry temperature of 220°C, sheath gas temperature of 400°C, mass range of 50-2000 *m/z*, and mobility (1/Ko) range of 0.55-1.90 V.s/cm^2^. Fragmentation data were acquired with a collision energy of 50 eV, with 2 PASEF (parallel accumulation serial fragmentation) MS/MS scans per cycle, with active exclusion release after 0.1 min, for a total cycle of 0.53 seconds.

After data acquisition, LC-MS/MS data was preprocessed using Bruker MetaboScape^®^ version 9.0.1 (Bruker-Daltonics, Billerica, MA, USA) using the MCube T-Rex 4D Metabolomics workflow for peak picking and alignment. The intensity threshold was set to 2000 counts after experimentally determining the baseline noise from bacterial samples and solvent blanks. Features were further processed using mpactR software^92^ with the following parameters: mispicked peak correction – ringing mass window of 0.5 atomic mass units (AMUs), isotopic mass window of 0.01 AMU with a maximum isotopic mass shift of 3 AMUs, and a t_R_ window of 0.05; in-source ion filtering threshold of 0.95 Spearman correlation; median coefficient of variation (CV) of technical replicates of 0.5; and 50% MeOH blank and unconditioned YPD media filtering at a 0.05 threshold. Features were log_10_ transformed to generate the heatmap in Fig 6C using MetaboAnalyst 6.0^93^. Annotations were performed using MetaboScape^®^ and NPAtlas^94^ databases with a 5 ppm threshold and were verified using the Competitive Fragmentation Modeling for Metabolite Identification (CFM-ID) spectra prediction^95^.

### MIC assays

Antifungal susceptibility testing was evaluated by broth microdilution MIC assay in flat-bottom 96-well plates. Assays were performed in a total volume of 200 uL per well, with 2-fold dilutions of each drug in YPD or *Ec*-YPD. Plates were incubated for 24 hours at 30°C before OD_600_ values were determined on a BioTek 800 TS Absorbance Reader. Each condition was tested in biological and technical duplicates. Growth was normalized to the no drug control.

## Acknowledgements

We thank the Martens lab for providing *Bacteroides* strains and germ-free fecal samples, thanks to the Sandkvist, Mobley, and Huffnagle labs for bacterial strains, and the Apostolides lab for use of their osmometer. Funding for this study was provided by National Institutes of Health grant R35GM147894 to TRO, University of Michigan Rackham Predoctoral Fellowship to F.A.D., National Institutes of Health NIAID T32 AI007528 to F.A.D.. Solid-state NMR analysis was supported by the National Institutes of Health (NIH) grant R01AI173270 to T.W. Metabolomics analyses were supported by start up funds to M.J.B. and a University of Michigan Rackham Graduate Student Research Grant to J.M.K. Tulane University School of Medicine Pilot Funding to S.E.R. NIAID DP2AI177927, CIFAR Global Azrieli Scholars Program, Crohn’s and Colitis Foundation CDA 884308, and R21AI188719 to K.S.O.

## Author contributions

Conceptualization: F.A.D and T.R.O Investigation: F.A.D, K.S, J.M.K, J.B, and K.S.O

Formal analysis: F.A.D, K.S, J.M.K, J.S, K.S.O, S.E.R, T.W, M.J.B, and T.R.O

Writing: F.A.D, K.S, J.M.K, K.O, S.E.R, T.W, M.J.B, and T.R.O

Reviewing and Editing: F.A.D, K.S, J.S, K.S.O, S.E.R, T.W, M.J.B, and T.R.O

Funding acquisition: T.R.O, F.A.D, S.E.R, K.S.O, M.J.B, T.W.

## Supplemental Figures and Figure Legends

**Supplemental Figure 1:** *E. coli* grows under co-culture conditions.

Growth curves of *E. coli* grown in LB or YPD for 24 hours at 30°C.

**Supplemental Figure 2:** *C. albicans* cells are viable after 24 hours of co-culture with *E. coli*.

Brightfield and fluorescent microscopy of *C. albicans* alone (top), *C. albicans* + *E. coli* (middle), or heat-killed *C. albicans* (bottom). Cells were stained with calcofluor white to mark total cells and propidium iodide to mark dead cells. 40x magnification.

**Supplemental Figure 3:** TEM preparation did not preserve the inner fungal cell wall.

Representative transmission electron micrographs depicting abnormal inner cell walls of *C. albicans* grown alone (left) or in co-culture with *E. coli* (right).

**Supplemental Figure 4:** Flow cytometry gating strategy and comparison of cell size.

**A)** Fungal cells were gated based on forward and side scatter to capture only fungal cells. Similar gating was performed on *C. albicans* in monoculture (left) or in co-culture with *E. coli* (right). **B)** Forward scatter histogram depicting differences in cell size when *C. albicans* is grown in monoculture (grey) or in co-culture with *E. coli* (pink).

**Supplemental Figure 5:** Relationship between cell size and cell wall contents.

**A, B, C)** Representative histogram of cell wall staining (left) for mannan **(A)**, exposed β-1,3-glucan **(B)**, or chitin **(C)**. Gray peaks represent *C. albicans* grown in monoculture. Flow cytometric dot plots comparing cell size with cell wall components during *C. albicans* monoculture or co-culture with *E. coli* (right).

**Supplemental Figure 6**: Human fecal IgA binding to *C. albicans* grown in monoculture or co-culture with *E. coli.* Significance determined by Wilcoxon test.

**Supplemental Figure 7:** Representative dynamic gradients of polysaccharides in the cel wall.

**A)** Cross polarization (CP) spectra representing rigid components. **B)** 2 s direct polarization (DP) spectra representing mobile components.

**Supplemental Figure 8:** Cell wall mannan content in different *Candida* species

**A)** Flow cytometric quantification of cell wall mannan for the indicated *Candida* species.

**Supplemental Figure 9:** Co-culture with diverse bacterial species under aerobic and anaerobic conditions

**A, B)** Representative brightfield and fluorescent microscopy images (left) and mannan histograms (right) of C. albicans grown alone or with indicated bacterial species under aerobic **(A)** or anaerobic **(B)** conditions. 40X magnification.

**Supplemental Figure 10:** *E. coli*-conditioned YPD delays growth of *C. albicans*. Growth curve of *C. albicans* grown in YPD or *Ec*-YPD for 24 hours.

**Supplemental Figure 11: Abundance patterns of shared features from metabolomics analysis** Shared features from LC-MS/MS of organic supernatants from the indicated bacterial species and *Ec*-YPD fractions. Fraction 4 was active while fractions 3 and 5 were not active.

**Supplemental Figure 12:** Minimum inhibitory concentrations

**A)** Minimum inhibitory concentration assay for *C. albicans* grown in YPD or *Ec*-YPD and treated with increasing concentrations of amphotericin B, caspofungin, and fluconazole.

**Supplemental Table 1: Fungal and bacterial strains used in this study Supplemental Table 2: RNAseq**

**Supplemental Table 3: Plasmids used in this study Supplemental Table 4: Oligonucleotides used in this study**

## References

1. Nash, A. K. et al. The gut mycobiome of the Human Microbiome Project healthy cohort. Microbiome 5, 153 (2017).

2. Lionakis, M. S. & Netea, M. G. Candida and host determinants of susceptibility to invasive candidiasis. PLoS Pathog. 9, e1003079 (2013).

3. Neville, B. A., d’Enfert, C. & Bougnoux, M.-E. Candida albicans commensalism in the gastrointestinal tract. FEMS Yeast Res. 15, (2015).

4. Puel, A. Human inborn errors of immunity underlying superficial or invasive candidiasis. Hum. Genet. 139, 1011–1022 (2020).

5. Fan, D. et al. Activation of HIF-1α and LL-37 by commensal bacteria inhibits Candida albicans colonization. Nat. Med. 21, 808–814 (2015).

6. d’Enfert, C., et al. The impact of the Fungus-Host-Microbiota interplay upon Candida albicans infections: current knowledge and new perspectives. FEMS Microbiol. Rev. (2020) doi:10.1093/femsre/fuaa060.

7. Hall, R. A. Adapting to change: interactions of Candida albicans with its environment. Future Microbiol. 12, 931–934 (2017).

8. Brown, A. J. P., Brown, G. D., Netea, M. G. & Gow, N. A. R. Metabolism impacts upon Candida immunogenicity and pathogenicity at multiple levels. Trends Microbiol. 22, 614–622 (2014).

9. Pradhan, A. et al. Hypoxia Promotes Immune Evasion by Triggering β-Glucan Masking on the Candida albicans Cell Surface via Mitochondrial and cAMP-Protein Kinase A Signaling. MBio 9, (2018).

10. Cottier, F. et al. Remasking of Candida albicans β-Glucan in Response to Environmental pH Is Regulated by Quorum Sensing. MBio 10, (2019).

11. Sherrington, S. L. et al. Adaptation of Candida albicans to environmental pH induces cell wall remodelling and enhances innate immune recognition. PLoS Pathog. 13, e1006403 (2017).

12. Ballou, E. R. et al. Lactate signalling regulates fungal β-glucan masking and immune evasion. Nat Microbiol 2, 16238 (2016).

13. Ene, I. V. et al. Host carbon sources modulate cell wall architecture, drug resistance and virulence in a fungal pathogen. Cell. Microbiol. 14, 1319–1335 (2012).

14. Ene, I. V. et al. Cell Wall Remodeling Enzymes Modulate Fungal Cell Wall Elasticity and Osmotic Stress Resistance. MBio 6, e00986 (2015).

15. Tripathi, A., Liverani, E., Tsygankov, A. Y. & Puri, S. Iron alters the cell wall composition and intracellular lactate to affect Candida albicans susceptibility to antifungals and host immune response. J. Biol. Chem. 295, 10032–10044 (2020).

16. Lenardon, M. D., Sood, P., Dorfmueller, H. C., Brown, A. J. P. & Gow, N. A. R. Scalar nanostructure of the Candida albicans cell wall; a molecular, cellular and ultrastructural analysis and interpretation. Cell Surf 6, 100047 (2020).

17. Brown, G. D. & Gordon, S. Immune recognition. A new receptor for beta-glucans. Nature 413, 36–37 (2001).

18. Taylor, P. R. et al. Dectin-1 is required for beta-glucan recognition and control of fungal infection. Nat. Immunol. 8, 31–38 (2007).

19. Graus, M. S. et al. Mannan Molecular Substructures Control Nanoscale Glucan Exposure in Candida. Cell Rep. 24, 2432–2442.e5 (2018).

20. Childers, D. S. et al. Epitope Shaving Promotes Fungal Immune Evasion. MBio 11, (2020).

21. Yang, M. et al. Control of β-glucan exposure by the endo-1,3-glucanase Eng1 in Candida albicans modulates virulence. PLoS Pathog. 18, e1010192 (2022).

22. Anderson, F. M. et al. Candida albicans selection for human commensalism results in substantial within-host diversity without decreasing fitness for invasive disease. PLoS Biol. 21, e3001822 (2023).

23. Tso, G. H. W. et al. Experimental evolution of a fungal pathogen into a gut symbiont. Science 362, 589– 595 (2018).

24. Böhm, L. et al. The yeast form of the fungus Candida albicans promotes persistence in the gut of gnotobiotic mice. PLoS Pathog. 13, e1006699 (2017).

25. McCrory, C., Lenardon, M. & Traven, A. Bacteria-derived short-chain fatty acids as potential regulators of fungal commensalism and pathogenesis. Trends Microbiol. (2024) doi:10.1016/j.tim.2024.04.004.

26. McCrory, C. et al. The short-chain fatty acid crotonate reduces invasive growth and immune escape of Candida albicans by regulating hyphal gene expression. MBio 14, e0260523 (2023).

27. Noverr, M. C. & Huffnagle, G. B. Regulation of Candida albicans morphogenesis by fatty acid metabolites. Infect. Immun. 72, 6206–6210 (2004).

28. Avelar, G. M. et al. Impact of changes at the Candida albicans cell surface upon immunogenicity and colonisation in the gastrointestinal tract. The Cell Surface 8, 100084 (2022).

29. Guinan, J., Wang, S., Hazbun, T. R., Yadav, H. & Thangamani, S. Antibiotic-induced decreases in the levels of microbial-derived short-chain fatty acids correlate with increased gastrointestinal colonization of Candida albicans. Sci. Rep. 9, 8872 (2019).

30. Strus, M. et al. The in vitro activity of vaginal Lactobacillus with probiotic properties against Candida. Infect. Dis. Obstet. Gynecol. 13, 69–75 (2005).

31. Graham, C. E., Cruz, M. R., Garsin, D. A. & Lorenz, M. C. Enterococcus faecalis bacteriocin EntV inhibits hyphal morphogenesis, biofilm formation, and virulence of Candida albicans. Proc. Natl. Acad. Sci. U. S. A. 114, 4507–4512 (2017).

32. Hogan, D. A., Vik, A. & Kolter, R. A Pseudomonas aeruginosa quorum-sensing molecule influences Candida albicans morphology. Mol. Microbiol. 54, 1212–1223 (2004).

33. Morales, D. K. et al. Antifungal mechanisms by which a novel Pseudomonas aeruginosa phenazine toxin kills Candida albicans in biofilms. Mol. Microbiol. 78, 1379–1392 (2010).

34. MacAlpine, J. et al. A small molecule produced by Lactobacillus species blocks Candida albicans filamentation by inhibiting a DYRK1-family kinase. Nat. Commun. 12, 6151 (2021).

35. Cuskin, F. et al. Human gut Bacteroidetes can utilize yeast mannan through a selfish mechanism. Nature 517, 165–169 (2015).

36. Temple, M. J. et al. A Bacteroidetes locus dedicated to fungal 1,6-β-glucan degradation: Unique substrate conformation drives specificity of the key endo-1,6-β-glucanase. J. Biol. Chem. 292, 10639– 10650 (2017).

37. Charlet, R., Bortolus, C., Sendid, B. & Jawhara, S. Bacteroides thetaiotaomicron and Lactobacillus johnsonii modulate intestinal inflammation and eliminate fungi via enzymatic hydrolysis of the fungal cell wall. Sci. Rep. 10, 11510 (2020).

38. de Assis, L. J. et al. Nature of β-1,3-Glucan-Exposing Features on Candida albicans Cell Wall and Their Modulation. MBio e0260522 (2022) doi:10.1128/mbio.02605-22.

39. Gantner, B. N., Simmons, R. M. & Underhill, D. M. Dectin-1 mediates macrophage recognition of Candida albicans yeast but not filaments. EMBO J. 24, 1277–1286 (2005).

40. Cabib, E. & Bowers, B. Chitin and yeast budding. J. Biol. Chem. 246, 152–159 (1971).

41. McGreal, E. P. et al. The carbohydrate-recognition domain of Dectin-2 is a C-type lectin with specificity for high mannose. Glycobiology 16, 422–430 (2006).

42. Saijo, S. et al. Dectin-2 recognition of alpha-mannans and induction of Th17 cell differentiation is essential for host defense against Candida albicans. Immunity 32, 681–691 (2010).

43. Ost, K. S. et al. Adaptive immunity induces mutualism between commensal eukaryotes. Nature (2021) doi:10.1038/s41586-021-03722-w.

44. Chakraborty, A. et al. A molecular vision of fungal cell wall organization by functional genomics and solid-state NMR. Nat. Commun. 12, 1–12 (2021).

45. Ghassemi, N. et al. Solid-state NMR investigations of extracellular matrixes and cell walls of algae, bacteria, fungi, and plants. Chem. Rev. 122, 10036–10086 (2022).

46. Cheng, Q. et al. Molecular architecture of chitin and chitosan-dominated cell walls in zygomycetous fungal pathogens by solid-state NMR. Nat. Commun. 15, 8295 (2024).

47. Dickwella Widanage, M. C., et al. Adaptative survival of Aspergillus fumigatus to echinocandins arises from cell wall remodeling beyond β-1,3-glucan synthesis inhibition. Nat. Commun. 15, 6382 (2024).

48. Lamon, G. et al. Solid-state NMR molecular snapshots of Aspergillus fumigatus cell wall architecture during a conidial morphotype transition. Proc. Natl. Acad. Sci. U. S. A. 120, e2212003120 (2023).

49. Dickwella Widanage, M. C.., et al. Distinct echinocandin responses of Candida albicans and Candida auris cell walls revealed by solid-state NMR. Nat. Commun. 16, 6295 (2025).

50. Hou, G., Yan, S., Trébosc, J., Amoureux, J.-P. & Polenova, T. Broadband homonuclear correlation spectroscopy driven by combined R2(n)(v) sequences under fast magic angle spinning for NMR structural analysis of organic and biological solids. J. Magn. Reson. 232, 18–30 (2013).

51. Hirakawa, M. P. et al. Genetic and phenotypic intra-species variation in Candida albicans. Genome Res. 25, 413–425 (2015).

52. Dunn, M. J., Fillinger, R. J., Anderson, L. M. & Anderson, M. Z. Automated quantification of Candida albicans biofilm-related phenotypes reveals additive contributions to biofilm production. NPJ Biofilms Microbiomes 6, 36 (2020).

53. Li, X., Yan, Z. & Xu, J. Quantitative variation of biofilms among strains in natural populations of Candida albicans. Microbiology 149, 353–362 (2003).

54. Huang, M. Y., Woolford, C. A., May, G., McManus, C. J. & Mitchell, A. P. Circuit diversification in a biofilm regulatory network. PLoS Pathog. 15, e1007787 (2019).

55. MacCallum, D. M. et al. Property differences among the four major Candida albicans strain clades. Eukaryot. Cell 8, 373–387 (2009).

56. Ropars, J. et al. Gene flow contributes to diversification of the major fungal pathogen Candida albicans. Nat. Commun. 9, 1–10 (2018).

57. Blattner, F. R. et al. The complete genome sequence of Escherichia coli K-12. Science 277, 1453–1462 (1997).

58. Mobley, H. L. et al. Pyelonephritogenic Escherichia coli and killing of cultured human renal proximal tubular epithelial cells: role of hemolysin in some strains. Infect. Immun. 58, 1281–1289 (1990).

59. Mulvey, M. A., Schilling, J. D. & Hultgren, S. J. Establishment of a persistent Escherichia coli reservoir during the acute phase of a bladder infection. Infect. Immun. 69, 4572–4579 (2001).

60. Bandara, H. M. H. N., Cheung, B. P. K., Watt, R. M., Jin, L. J. & Samaranayake, L. P. Secretory products of Escherichia coli biofilm modulate Candida biofilm formation and hyphal development. J. Investig. Clin. Dent. 4, 186–199 (2013).

61. Matsuda, Y., Cho, O., Sugita, T., Ogishima, D. & Takeda, S. Culture Supernatants of Lactobacillus gasseri and L. crispatus Inhibit Candida albicans Biofilm Formation and Adhesion to HeLa Cells. Mycopathologia 183, 691–700 (2018).

62. Das, S. & Konwar, B. K. Inhibiting pathogenicity of vaginal Candida albicans by lactic acid bacteria and MS analysis of their extracellular compounds. APMIS (2024) doi:10.1111/apm.13365.

63. Homann, O. R., Dea, J., Noble, S. M. & Johnson, A. D. A phenotypic profile of the Candida albicans regulatory network. PLoS Genet. 5, e1000783 (2009).

64. Noble, S. M., French, S., Kohn, L. A., Chen, V. & Johnson, A. D. Systematic screens of a Candida albicans homozygous deletion library decouple morphogenetic switching and pathogenicity. Nat. Genet. 42, 590–598 (2010).

65. Román, E., Nombela, C. & Pla, J. The Sho1 adaptor protein links oxidative stress to morphogenesis and cell wall biosynthesis in the fungal pathogen Candida albicans. Mol. Cell. Biol. 25, 10611–10627 (2005).

66. Nagahashi, S. et al. Isolation of CaSLN1 and CaNIK1, the genes for osmosensing histidine kinase homologues, from the pathogenic fungus Candida albicans. Microbiology 144 (Pt 2), 425–432 (1998).

67. Sellam, A. et al. The p38/HOG stress-activated protein kinase network couples growth to division in Candida albicans. PLoS Genet. 15, e1008052 (2019).

68. Plaine, A. et al. Functional analysis of Candida albicans GPI-anchored proteins: roles in cell wall integrity and caspofungin sensitivity. Fungal Genet. Biol. 45, 1404–1414 (2008).

69. Lee, K. K. et al. Elevated cell wall chitin in Candida albicans confers echinocandin resistance in vivo. Antimicrob. Agents Chemother. 56, 208–217 (2012).

70. Avelar, G. M. et al. A CO2 sensing module modulates β-1,3-glucan exposure in Candida albicans. MBio e0189823 (2024) doi:10.1128/mbio.01898-23.

71. Zhang, M. et al. CO2 potentiates echinocandin efficacy during invasive candidiasis therapy via dephosphorylation of Hsp90 by Ptc2 in condensates. Proc. Natl. Acad. Sci. U. S. A. 122, e2417721122 (2025).

72. Gutierrez, D. et al. Antibiotic-induced gut metabolome and microbiome alterations increase the susceptibility to Candida albicans colonization in the gastrointestinal tract. FEMS Microbiol. Ecol. 96, (2020).

73. Savage, H. P. et al. Epithelial hypoxia maintains colonization resistance against Candida albicans. Cell Host Microbe (2024) doi:10.1016/j.chom.2024.05.008.

74. Heredia, M. Y., Gunasekaran, D., Ikeh, M. A. C., Nobile, C. J. & Rauceo, J. M. Transcriptional regulation of the caspofungin-induced cell wall damage response in Candida albicans. Curr. Genet. 66, 1059–1068 (2020).

75. Rodríguez-Peña, J. M., García, R., Nombela, C. & Arroyo, J. The high-osmolarity glycerol (HOG) and cell wall integrity (CWI) signalling pathways interplay: a yeast dialogue between MAPK routes. Yeast 27, 495–502 (2010).

76. Brown, A. J. P. et al. Stress adaptation in a pathogenic fungus. J. Exp. Biol. 217, 144–155 (2014).

77. Page, W. J. & Tigerstrom, M. V. Aminochelin, a Catecholamine Siderophore Produced by Azotobacter vinelandii. Microbiology 134, 453–460 (1988).

78. Baars, O., Zhang, X., Morel, F. M. M. & Seyedsayamdost, M. R. The siderophore metabolome of Azotobacter vinelandii. Appl. Environ. Microbiol. 82, 27–39 (2015).

79. Shi, J. et al. Discovery and biosynthesis of guanipiperazine from a NRPS-like pathway. Chem. Sci. 12, 2925–2930 (2021).

80. 焦瑞华戈惠明 徐响 樊瑞. Guanine piperazine compound and its preparation method and application. Patent (2021).

81. Forseth, R. R. et al. Homologous NRPS-like gene clusters mediate redundant small-molecule biosynthesis in Aspergillus flavus. Angew. Chem. Int. Ed Engl. 52, 1590–1594 (2013).

82. Liu, L. et al. Asperorydines A-M: Prenylated tryptophan-derived alkaloids with neurotrophic effects from Aspergillus oryzae. J. Org. Chem. 83, 812–822 (2018).

83. Xiang, Y. et al. Asperorydines N-P, three new cyclopiazonic acid alkaloids from the marine-derived fungus Aspergillus flavus SCSIO F025. Fitoterapia 150, 104839 (2021).

84. Noble, S. M. & Johnson, A. D. Strains and strategies for large-scale gene deletion studies of the diploid human fungal pathogen Candida albicans. Eukaryot. Cell 4, 298–309 (2005).

85. Lesage, A., Bardet, M. & Emsley, L. Through-bond Carbon−Carbon connectivities in disordered solids by NMR. J. Am. Chem. Soc. 121, 10987–10993 (1999).

86. Lee, D. W., Hong, C. P. & Kang, H. A. An effective and rapid method for RNA preparation from non-conventional yeast species. Anal. Biochem. 586, 113408 (2019).

87. Aranda, P. S., LaJoie, D. M. & Jorcyk, C. L. Bleach gel: a simple agarose gel for analyzing RNA quality. Electrophoresis 33, 366–369 (2012).

88. Bolger, A. M., Lohse, M. & Usadel, B. Trimmomatic: a flexible trimmer for Illumina sequence data. Bioinformatics 30, 2114–2120 (2014).

89. Bray, N. L., Pimentel, H., Melsted, P. & Pachter, L. Near-optimal probabilistic RNA-seq quantification. Nat. Biotechnol. 34, 525–527 (2016).

90. Love, M. I., Huber, W. & Anders, S. Moderated estimation of fold change and dispersion for RNA-seq data with DESeq2. Genome Biol. 15, 550 (2014).

91. Kolberg, L. et al. g:Profiler-interoperable web service for functional enrichment analysis and gene identifier mapping (2023 update). Nucleic Acids Res. 51, W207–W212 (2023).

92. Mason, A. R. et al. mpactR: an R adaptation of the metabolomics peak analysis computational tool (MPACT) for use in reproducible data analysis pipelines. Microbiol. Resour. Announc. 14, e0099724 (2025).

93. Pang, Z. et al. MetaboAnalyst 6.0: towards a unified platform for metabolomics data processing, analysis and interpretation. Nucleic Acids Res. 52, W398–W406 (2024).

94. van Santen, J. A. et al. The Natural Products Atlas: An open access knowledge base for microbial natural products discovery. ACS Cent. Sci. 5, 1824–1833 (2019).

95. Allen, F., Pon, A., Wilson, M., Greiner, R. & Wishart, D. CFM-ID: a web server for annotation, spectrum prediction and metabolite identification from tandem mass spectra. Nucleic Acids Res. 42, W94–9 (2014).

