## Supplementary figures and images for "Bacterial metabolites induce cell wall remodeling, antifungal resistance, and immune recognition of commensal fungi"

### FigS1

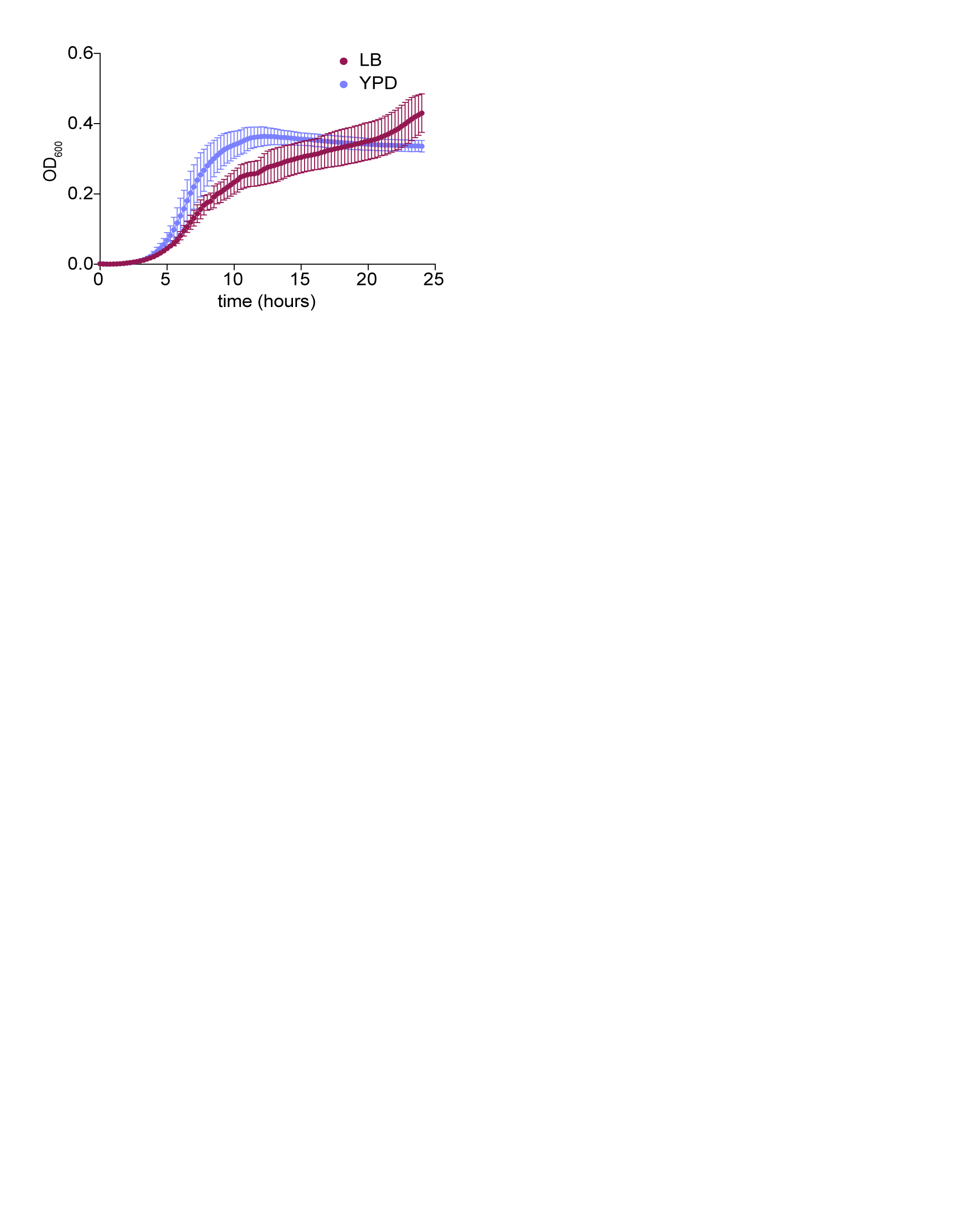

### FigS2

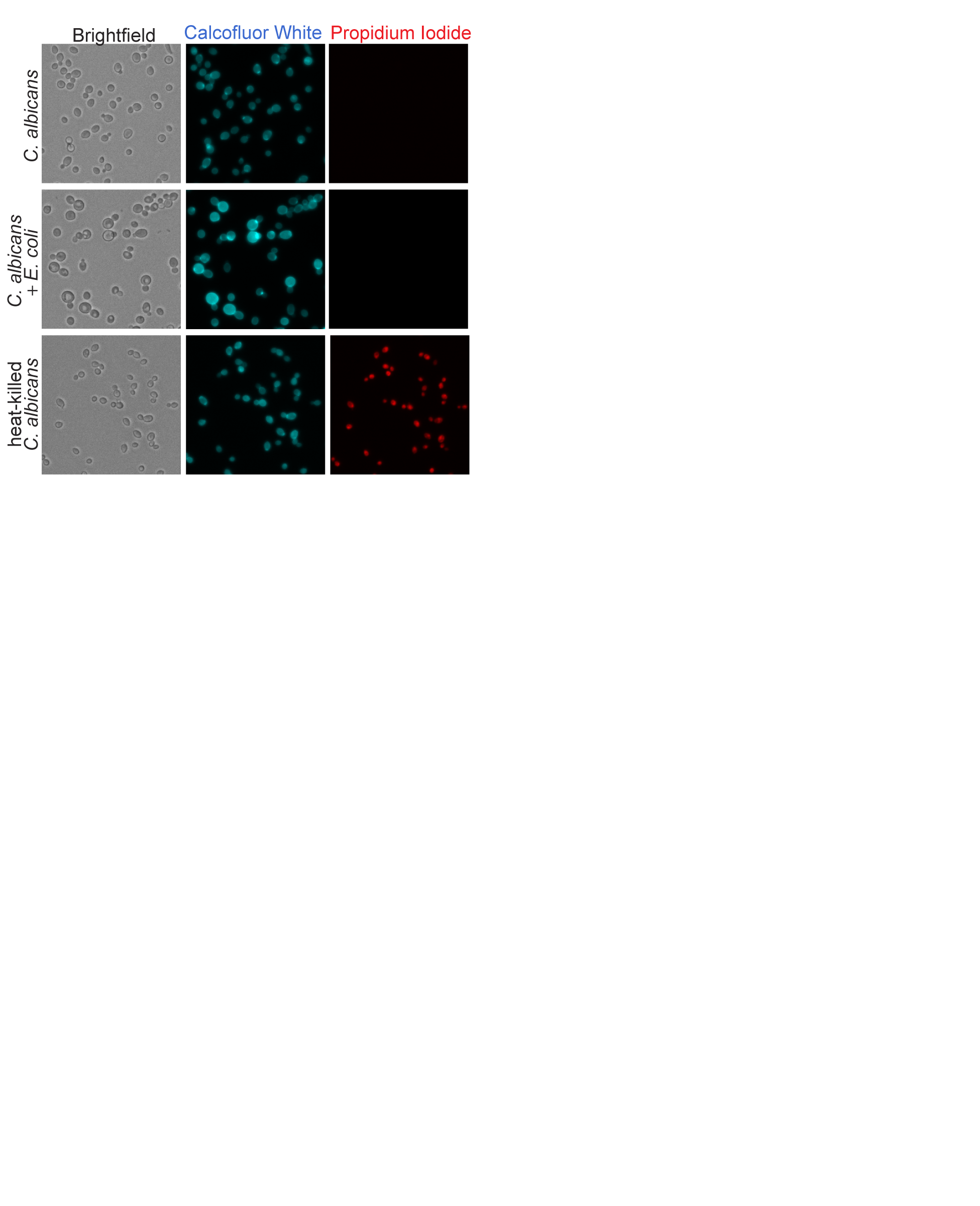

### FigS3

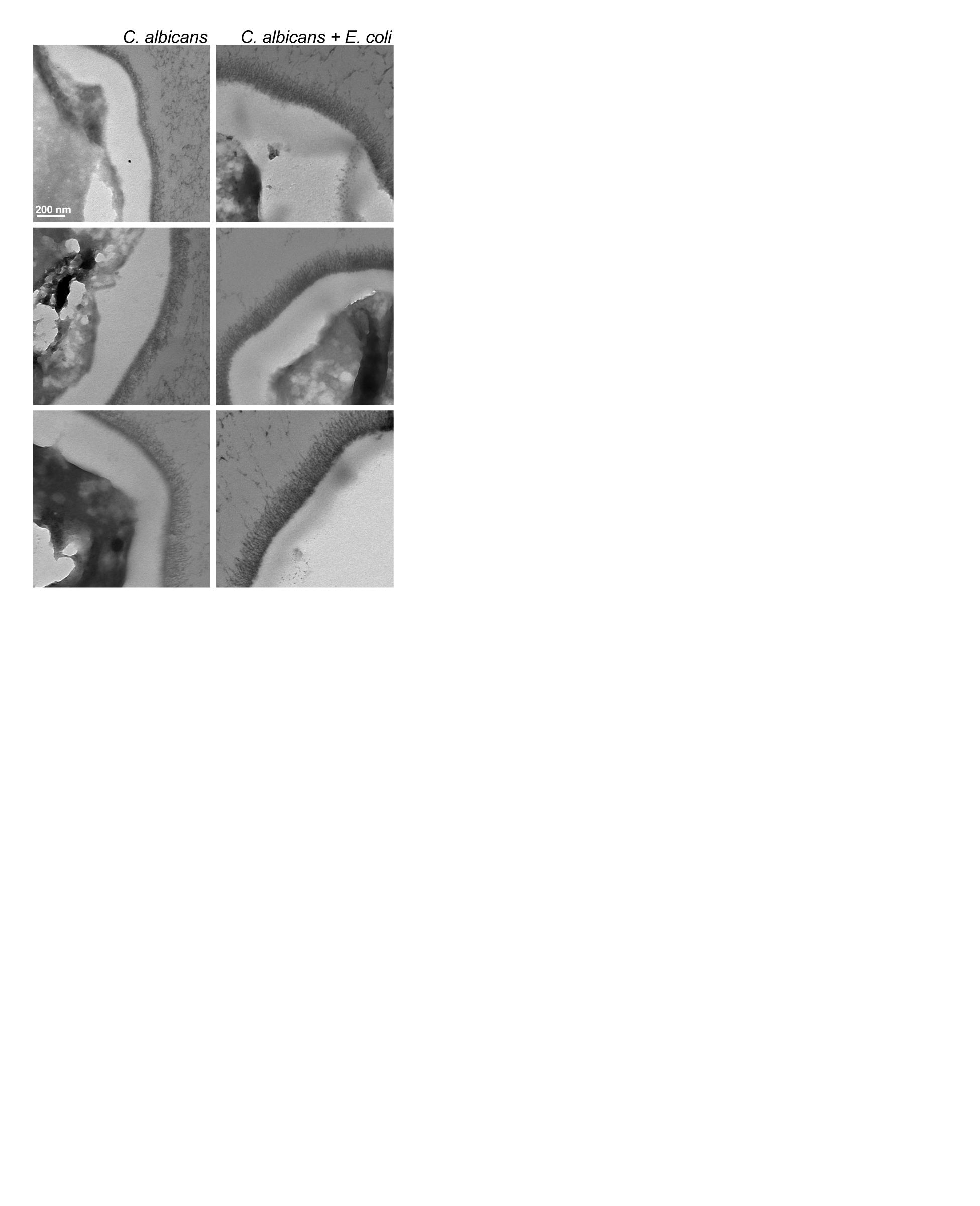

### FigS4

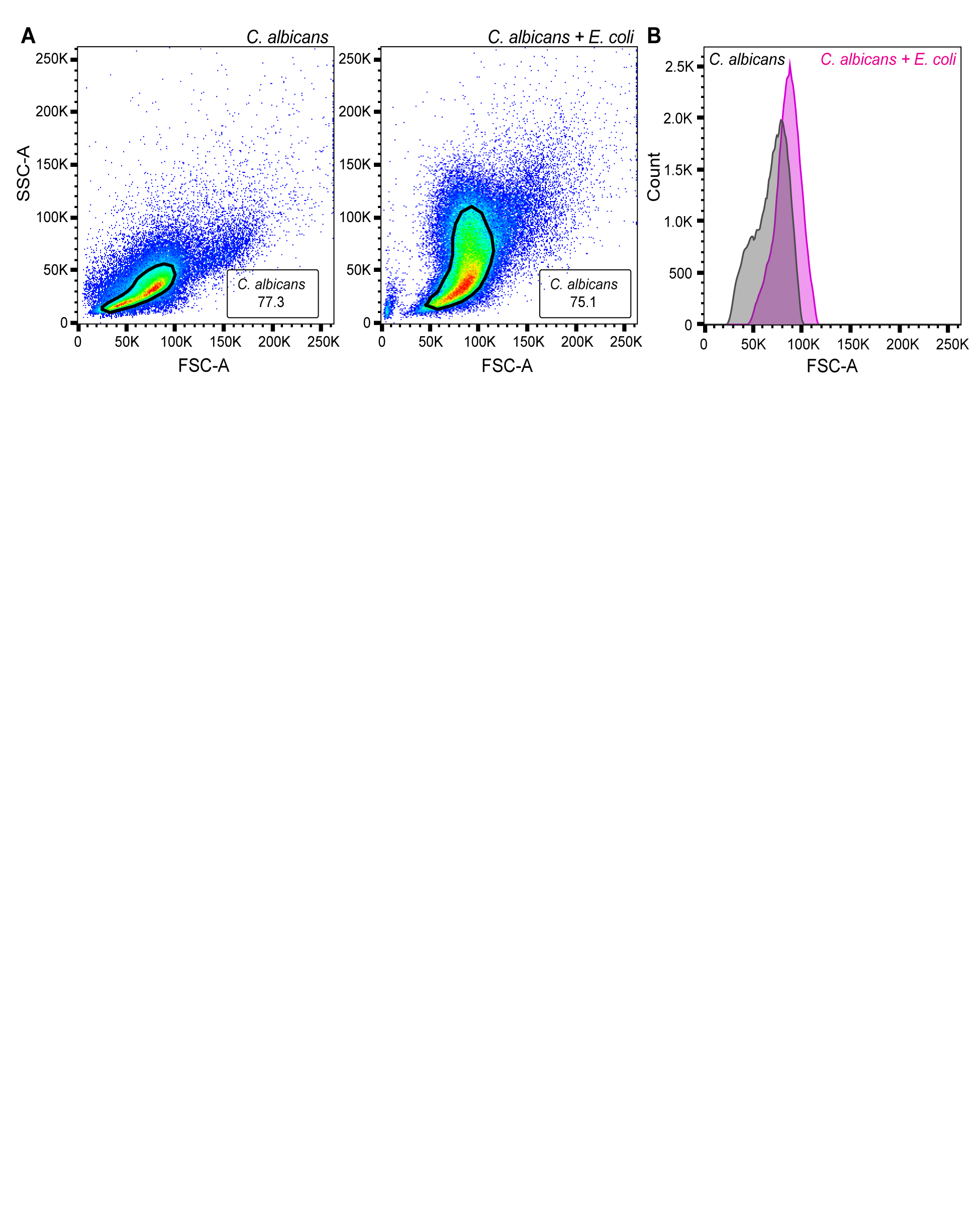

### FigS5

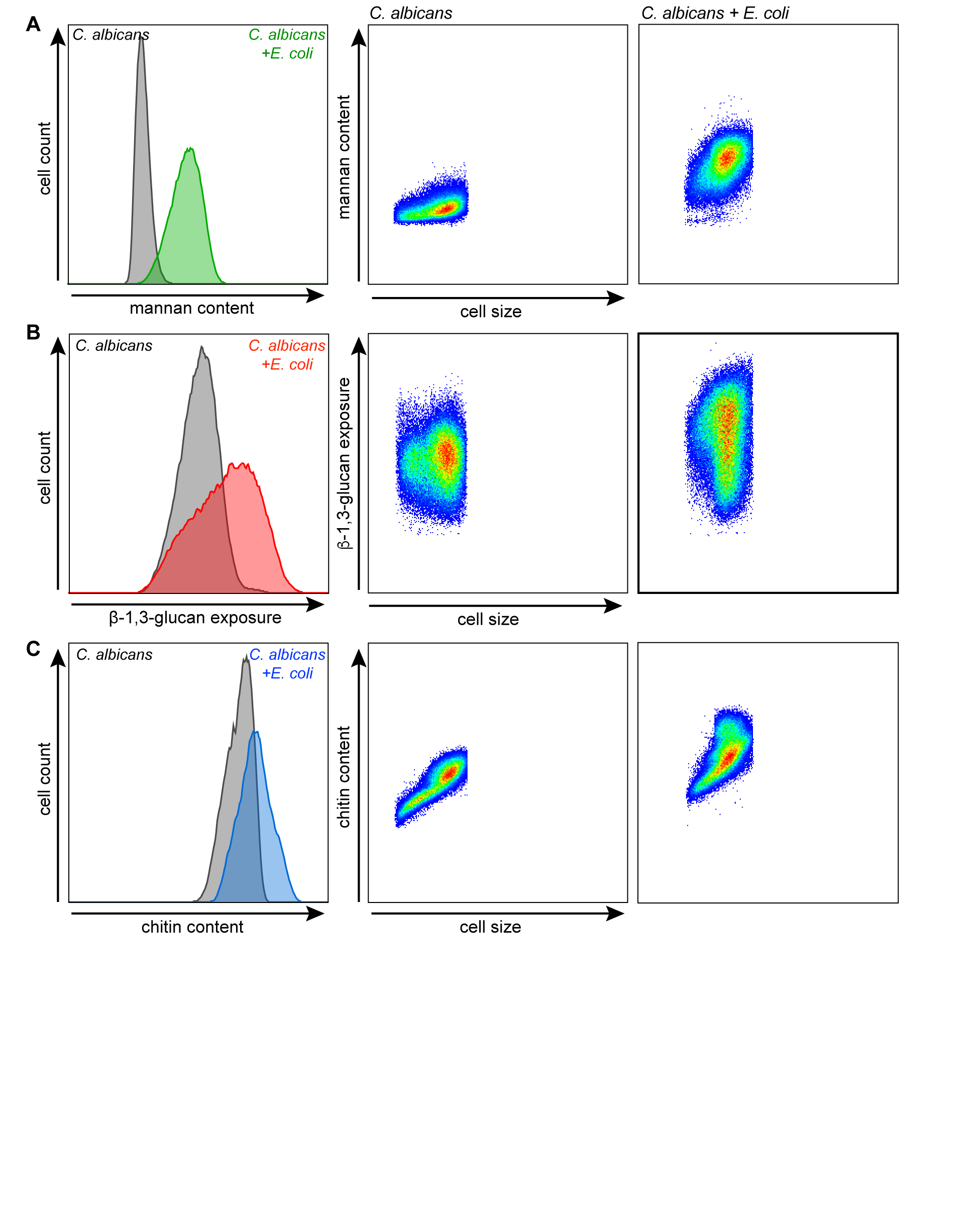

### FigS6

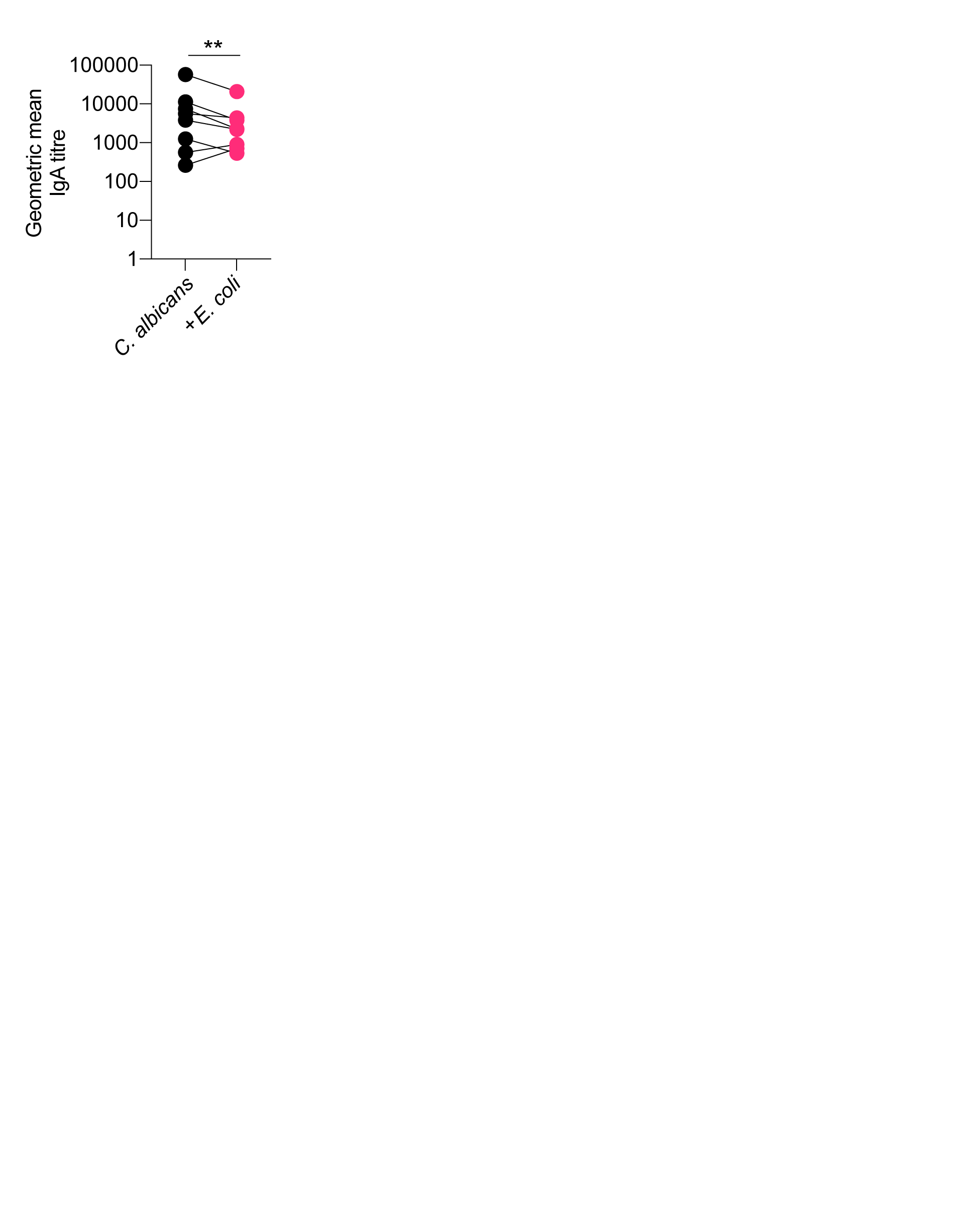

### FigS7

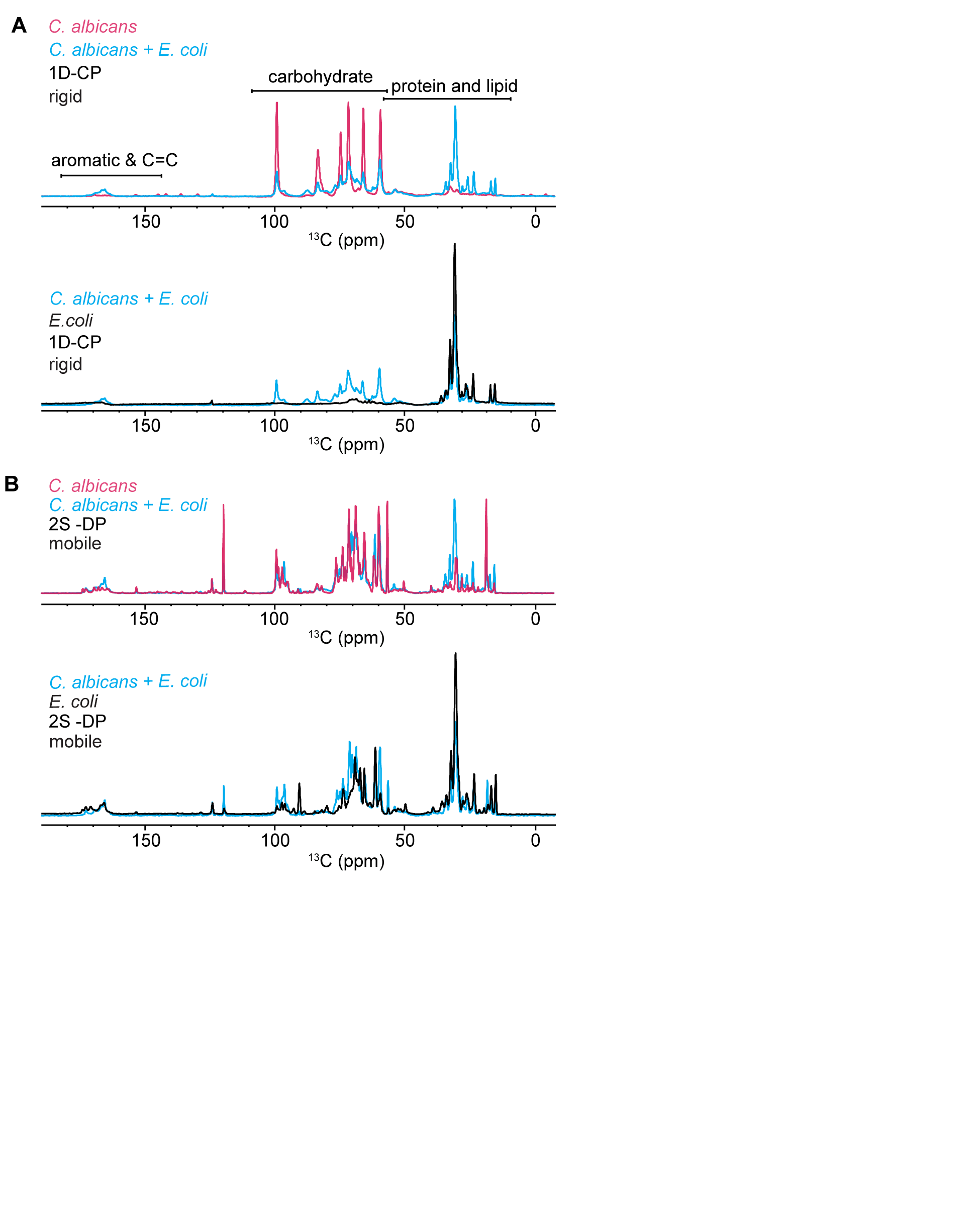

### FigS8

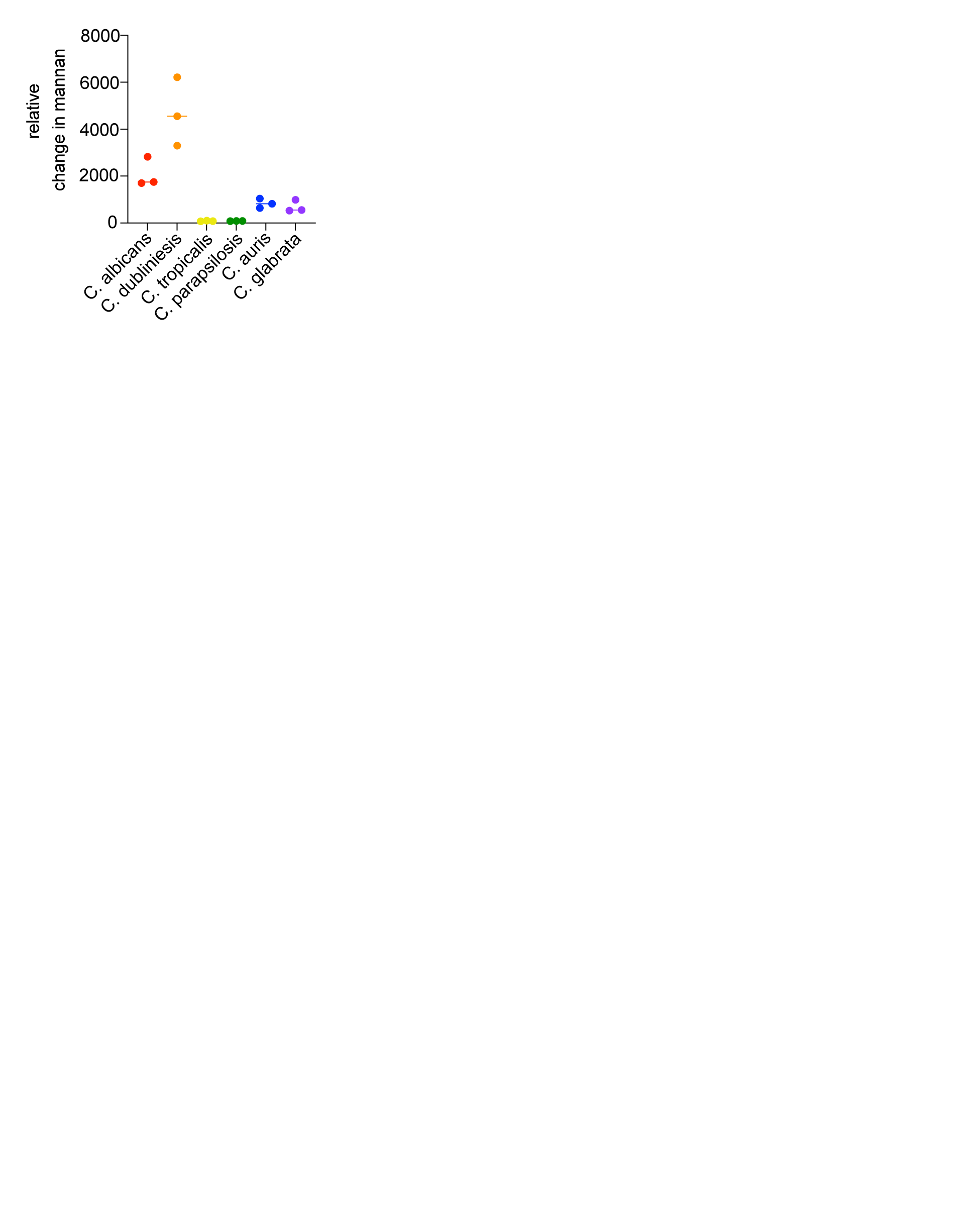

### FigS9

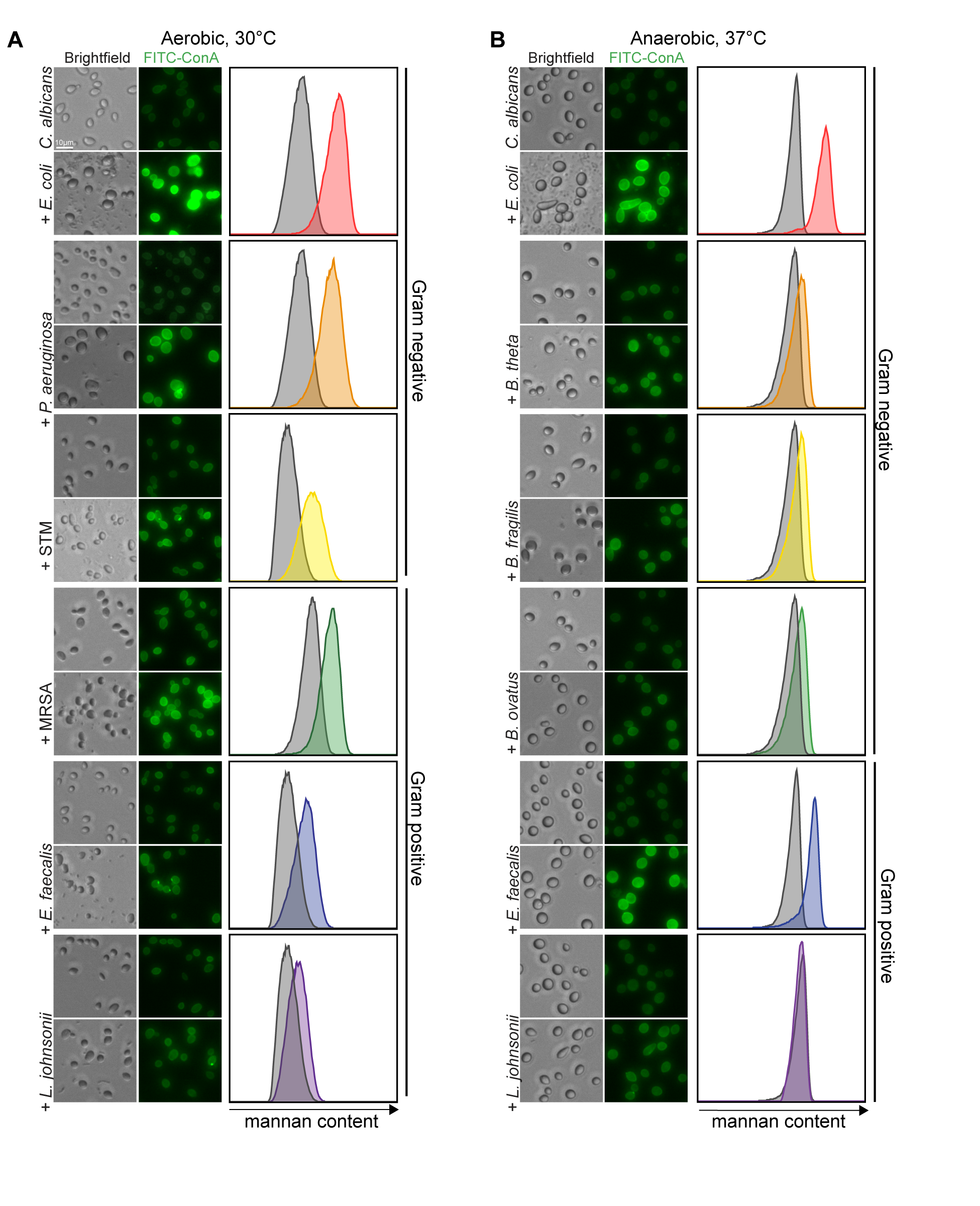

### FigS10

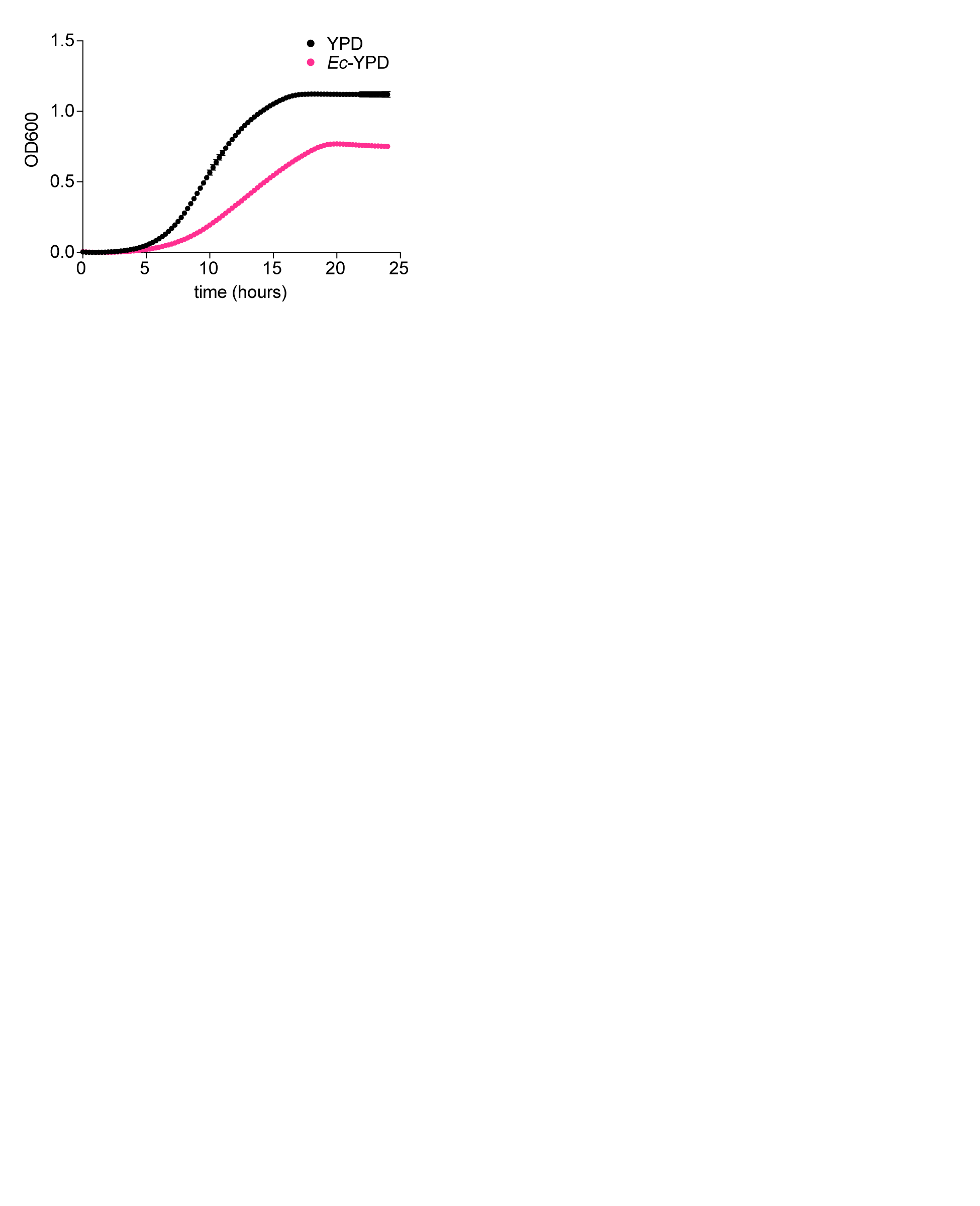

### FigS11

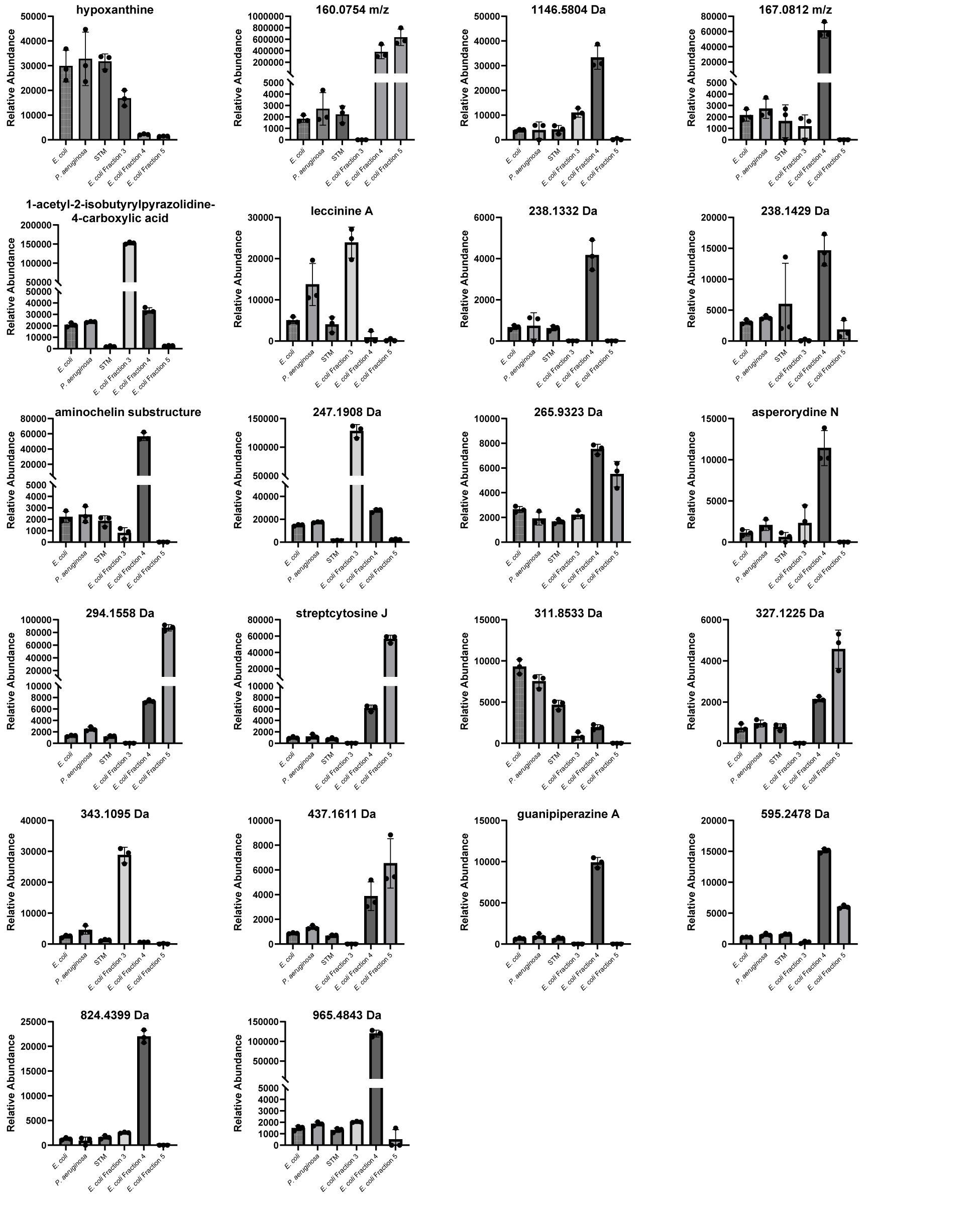

### FigS12

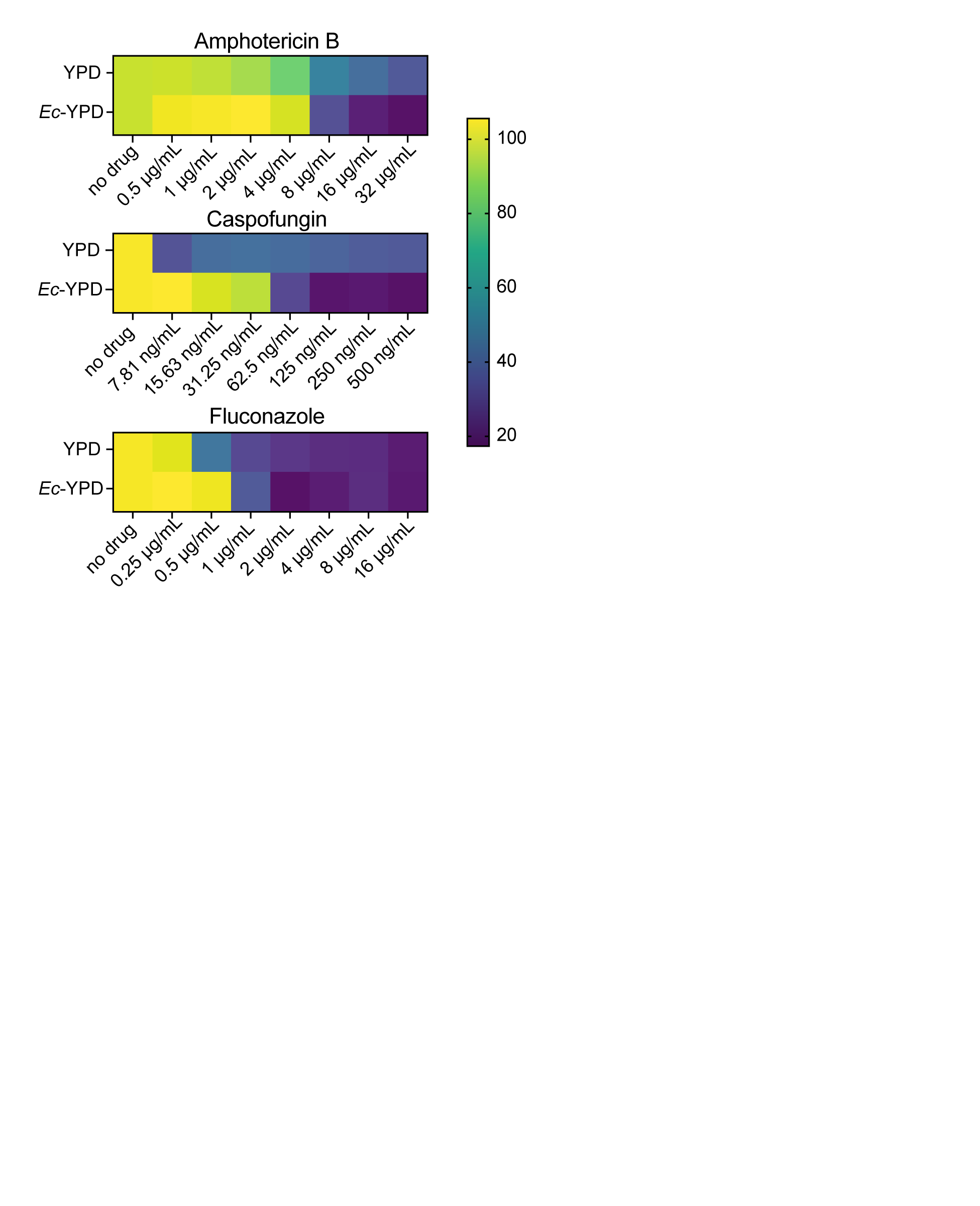
